# A Central Role for Canonical PRC1 in Shaping the 3D Nuclear Landscape

**DOI:** 10.1101/2019.12.15.876771

**Authors:** Shelagh Boyle, Ilya M. Flyamer, Iain Williamson, Dipta Sengupta, Wendy A. Bickmore, Robert S. Illingworth

## Abstract

Polycomb group (PcG) proteins silence gene expression by chemically and physically modifying chromatin. A subset of PcG target loci are compacted and cluster in the nucleus to form observable bodies; a conformation which is thought to contribute to gene silencing. However, how these interactions influence gross nuclear organisation and their relationship with transcription remains poorly understood. Here we examine the role of Polycomb Repressive Complex 1 (PRC1) in shaping 3D genome organization in mouse embryonic stem cells (mESCs). Using a combination of imaging and Hi-C analyses we show that PRC1-mediated long-range interactions are independent of CTCF and can bridge sites at a megabase scale. Impairment of PRC1 enzymatic activity does not directly disrupt these interactions. We demonstrate that PcG targets coalesce in vivo, and that developmentally induced expression of one of the target loci disrupts this spatial arrangement. Finally, we show that transcriptional activation and the loss of PRC1-mediated interactions are seperable events. These findings provide important insights into the function of PRC1, whilst highlighting the complexity of this regulatory system.

**Highlights:**

1. Loss of RING1B substantially disrupts nuclear architecture.
2. PRC1 mediated looping can occur at a Mb scale and is independent of CTCF.
3. Polycomb mediated looping is driven by canonical PRC1 complexes.
4. Multimeric PRC1-mediated interactions occur in vitro and in vivo.
5. Disruption of PRC1-mediated looping is independent of gene activation.

## Introduction

The spatial and temporal fidelity of development is controlled by transcription factors that act in concert with the epigenome to regulate gene expression programmes (Simon and Kingston 2013; Atlasi and Stunnenberg 2017). Polycomb repressive complexes (PRCs), a family of essential epigenetic regulators, modify chromatin to propagate a repressed but transcriptionally poised state (Brookes and Pombo 2009; Simon and Kingston 2013; Voigt et al. 2013; Blackledge et al. 2015; Schuettengruber et al. 2017). PRC1 and 2, the two principal members of this family, prevent unscheduled differentiation by targeting and restricting the expression of genes encoding key developmental regulators. The deposition of H2AK119ub1 and H3K27me1/2/3 by RING1A/B (PRC1) and EZH1/2 (PRC2) respectively, is required for the efficient placement of PRCs at target loci (Poux et al. 2001; Cao et al. 2002; Czermin et al. 2002; Kuzmichev et al. 2002; Muller et al. 2002; de Napoles et al. 2004; Wang et al. 2004a; Wang et al. 2004b; Margueron et al. 2009; Blackledge et al. 2014; Cooper et al. 2014; Kalb et al. 2014).

The core of PRC1 comprises a heterodimer of RING1A/B and one of six PCGF RING finger proteins. Deposition of H2AK119Ub is driven primarily by variant PRC1s (vPRC1 - RING1A/B complexed with either of PCGF1, 3, 5 or 6) which have enhanced E3-ligase activity due to an association with either RYBP or YAF2 (Rose et al. 2016; Fursova et al. 2019). Combinatorial deletion of PCGF1, 3, 5 and 6 in mouse embryonic stem cells (mESCs) leads to substantial gene misregulation and highlights the importance of vPRC1s in transcriptional control (Fursova et al. 2019). In contrast, canonical PRC1 (cPRC1; core heterodimer of RING1A or B with PCGF2 or 4) has lower catalytic activity and is instead associated with subunits that alter chromatin structure and topology (Francis et al. 2004; Grau et al. 2011; Isono et al. 2013; Blackledge et al. 2014; Taherbhoy et al. 2015; Wani et al. 2016; Kundu et al. 2017; Lau et al. 2017; Plys et al. 2019; Tatavosian et al. 2019). In line with this function, a subset of PRC1 targets fold into short discrete self-interacting domains (20-140 kb), exemplified by the conformation of the transcriptionally-silent *Hox* clusters in mESCs (Eskeland et al. 2010; Williamson et al. 2014; Vieux-Rochas et al. 2015; Kundu et al. 2017). However unlike topologically associated domains (TADs), which are somewhat structurally invariant across different cells types, PRC1 mediated domains are developmentally dynamic and are eroded upon gene activation and the loss of PRC1 association (Lieberman-Aiden et al. 2009; Eskeland et al. 2010; Dixon et al. 2012; Nora et al. 2012; Williamson et al. 2012; Rao et al. 2014; Bonev et al. 2017; Kundu et al. 2017). In addition to local chromatin folding, PRC1 coordinates interactions between distally located target sites (Isono et al. 2013; Schoenfelder et al. 2015; Kundu et al. 2017). Consequently, genomic loci which are separated by large distances in the linear genome can be brought into close spatial proximity. In Drosophila, this juxtaposition has been suggested to enhance transcriptional repression, but direct evidence for this in mammalian cells is lacking (Bantignies et al. 2003; Bantignies et al. 2011; Eagen et al. 2017; Ogiyama et al. 2018).

CBX and PHC subunits are thought to be the components of PRC1 which are primarily responsible for mediating these chromatin structures. CBX2, 6 and 8, mammalian homologues of Drosophila Polycomb, contain a positively charged intrinsically disordered region (IDR) that can compact nucleosomal arrays in vitro (Grau et al. 2011). Neutralising amino-acid substitutions in the IDR of CBX2 lead to some loss of PRC1-mediated gene repression and axial patterning defects in mice (Lau et al. 2017). Polyhomoeotic (PHC) proteins can make both homo- and hetero-meric head-to-tail interactions via their sterile alpha motif (SAM) domain (Isono et al. 2013), allowing multiple cPRC1s to oligomerise and thus to physically connect regions of the genome. Disruption of the SAM domain ablates these interactions leading to the loss of both local interaction domains and PRC1 mediated looping (Kundu et al. 2017) and resulting in gene de-repression and skeletal abnormalities in mice (Isono et al. 2013). Furthermore, loss of these architectural PRC1 subunits leads to the dissolution of nanometre scale ‘polycomb bodies’ containing high local concentrations of polycomb proteins (Isono et al. 2013; Wani et al. 2016; Plys et al. 2019; Tatavosian et al. 2019). These data support the idea that CBX and PHC proteins bestow cPRC1 with the capacity to fold chromatin into discrete nuclear domains and suggest a mechanistic role for chromatin interactions and nuclear clustering in PRC1-mediated transcriptional repression.

However, this emerging view raises some important questions. What factors determine which distal PRC1 targets will physically interact? Does PRC1 create a topology that anchors multiple loci simultaneously in a single cell and, if so, do such structures occur in vivo? What is the cause/consequence relationship between chromatin structure and gene de-repression in cells lacking RING1B? In this study, we employ both Hi-C and DNA Fluorescence in Situ Hybridisation (FISH) in mESCs and embryonic mouse tissue to investigate how PRC1 contributes to nuclear organisation. We find that PRC1 has a substantial effect on chromosomal architecture that is disproportionate to the fraction of the genome it occupies. These structures rely on canonical PRC1, are independent of CTCF and persist even when RING1B catalytic activity is substantially impaired. Our findings provide key insights into the manner in which PRC1 dictates the 3D topology of the mammalian genome.

## Results

### Loss of RING1B Disrupts Nuclear Clustering of Polycomb Targets

RING1B is the primary RING1 homolog expressed in mESCs and in its absence levels of the PRC1 complex are substantially reduced (Leeb and Wutz 2007; Endoh et al. 2008; Eskeland et al. 2010). DAPI staining of 2D nuclear preparations revealed a significant increase in nuclear area in the absence of PRC1 (*Ring1b^-/-^*) when compared to parental *Ring1b^+/+^* mESCs (**Fig. 1A**; Supplemental Fig. S1A). The polycomb system promotes cell proliferation, in part, by negatively regulating inhibitors of the cell cycle (Jacobs et al. 1999; Gil et al. 2004; Bracken et al. 2007; Chen et al. 2009). However, fluorescence-activated cell sorting (FACS) did not identify an altered cell cycle profile between *Ring1b^+/+^* and *Ring1b^-/-^* cells (Supplemental Fig. S1B). This suggested that the increase in nuclear size in *Ring1b^-/-^* cells is a direct consequence of PRC1 depletion on nuclear structure, frather than an accumulation of cells in G2.

**Figure 1.**
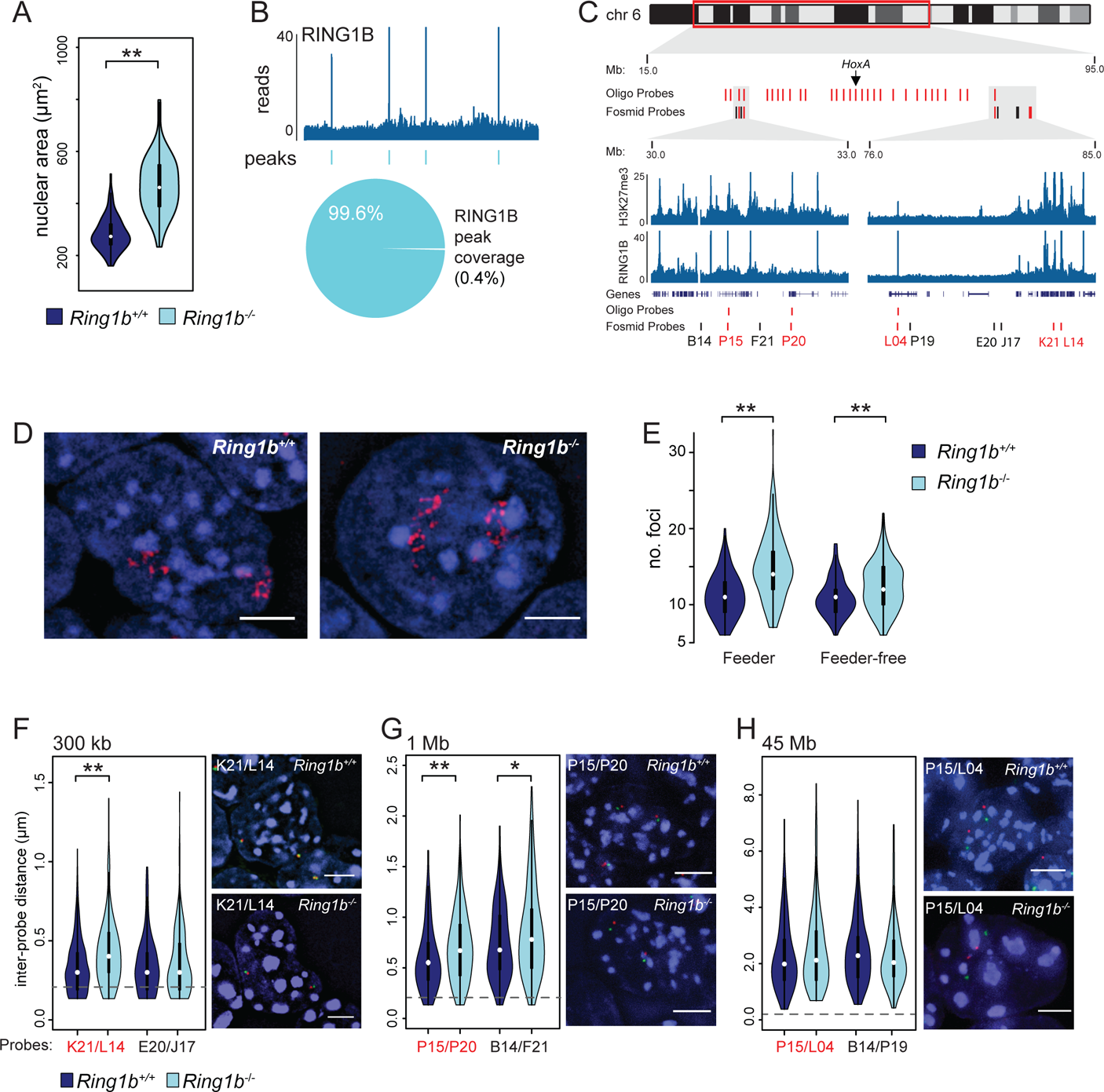
PRC1 dependent nuclear clustering of polycomb targets. (**A**) Violin plot depicting the nuclear area (μm^2^) of *Ring1b^+/+^* and *Ring1b^-/-^*mESCs determined by DAPI staining of 2D preparations. **p = 9.25×10^-23^; Mann whitney test. (**B**) Example ChIP-seq profile and called peaks for RING1B in wildtype mESCs (Chr 6: 62.5 - 68.5 Mb - mm9 genome assembly)(Illingworth et al. 2015). The pie chart below shows the sum coverage of all RING1B peaks as a fraction of the uniquely mappable portion of the mouse genome. (**C**) Ideogram of chromosome 6 showing the location of oligonucleotide and fosmid probes used in panels (**D-H**) and zoomed browser tracks of RING1B and H3K27me3 ChIP-seq from wildtype mESCs (Illingworth et al. 2015). Genome co-ordinates from the mm9 genome assembly. (**D**) Representative 3D FISH image of the chromosome 6 polycomb positive oligonucleotide probe signal in the nuclei of *Ring1b^+/+^* and *Ring1b^-/-^* mESCs (scale bar = 5 μm). (**E**) Violin plot depicting the number of discrete foci in *Ring1b^+/+^* and *Ring1b^-/-^* mESCs detected by FISH with the chromosome 6 polycomb positive oligonucleotide probe. Two independent *Ring1b^-/-^* clones and their associated wildtype parental mESCs are indicated (‘feeder’ and ‘feeder-free’; (Leeb and Wutz 2007; Illingworth et al. 2015)). **p = 9.07×10^-17^ and **5.33×10^-6^ for feeder and feeder free respectively; Mann Whitney test. (**F-H**) Violin plots of inter-probe distances (μm) for the indicated fosmids (locations shown in (**C**) with representative images for both *Ring1b^+/+^* and *Ring1b^-/-^*mESCs (scale bar = 5 μm)). Probes separated by less than 0.2 μm (dashed grey line) are considered to be co-localised. **p = 4.23×10^-07^ (**F**) and **p = 9.47×10^-04^ and *3.47×10^-02^ (**G**); Mann Whitney test.

Canonical PRC1 can directly alter chromatin structure, and ensemble chromatin conformation capture assays (e.g. Hi-C) have demonstrated that polycomb target sites can physically interact and so affect nuclear organisation (Schoenfelder et al. 2015; Vieux-Rochas et al. 2015; Bonev et al. 2017; Kundu et al. 2017) (Joshi et al. 2015). However, analysis of ChIP-seq data indicates that only a very small fraction (0.4%) of the mESC genome has pronounced RING1B occupancy (**Fig. 1B**; (Illingworth et al. 2015)). How then could the loss of RING1B/PRC1 at discrete sites lead to such a profound impact on nuclear size? To address this question, we determined the spatial arrangement of polycomb (PcG) target loci within individual nuclei and how this was disrupted in cells lacking RING1B. For this we needed a suitable 3D-FISH probe that would simultaneously detect multiple PcG target loci. We therefore designed a custom fluorescently labelled oligonucleotide pool covering 30 non-contiguous 20 kb windows along chromosome 6, each of which was centred on an individual PcG target locus (H3K27me3 positive peaks from ChIP-seq data; (Illingworth et al. 2015); **Fig. 1C**; Supplemental Table S1). Hybridisation to metaphase spreads confirmed the specificity and efficiency of the oligonucleotide probe (Supplemental Fig. 1C). In interphase, quantitation of foci by blind scoring demonstrated a marked co-localisation of polycomb targets. A median score of six foci per chromosome suggested that at least five or more PcG target sites might simultaneously localise to a single focus (Fig. 1D, E). The mean number of foci detected was significantly increased in each of two independent *Ring1b*^-/-^ mESC lines (p = 9.1×10^-17^ and 5.3×10^-6^ for feeder dependent and feeder independent ESC lines respectively (Leeb and Wutz 2007; Illingworth et al. 2015)(**Fig. 1E**), suggesting that polycomb targets cluster together in a PRC1 dependent manner in mESCs.

It has been shown that polycomb-mediated interactions can occur over large genomic distances (>10 Mb), (Joshi et al. 2015; Schoenfelder et al. 2015; Vieux-Rochas et al. 2015; Bonev et al. 2017; Kundu et al. 2017). To determine if polycomb site clustering was restricted by the linear proximity of target sites along the genome we performed 3D-FISH using fluorescently labelled fosmid probes targeting polycomb positive (PcG+) and negative (PcG-) loci (presence or absence of H3K27me3 respectively) located along the same region of chromosome 6 (**Fig. 1C**). PcG+ sites relatively close to each other in the linear genome (300 kb and 1 Mb) showed a significant increase in inter-probe distances in cells lacking RING1B (Fig. 1F, G and Supplemental Fig. 1D, E). In contrast, loss of RING1B did not significantly impact on the inter-probe distances when the probes were separated by 45 Mb (**Fig. 1H** and Supplemental Fig. 1F). These findings suggest that although long-range sites can be detected in close proximity within a population of cells, PRC1-mediated associations are generally constrained by chromosome topology and favour more proximal contacts. Interestingly, PcG-‘control’ probes separated by 1 Mb also showed a significant increase in inter-probe distance in *Ring1b^-/-^* mESCs despite lacking detectible H3K27me3 or RING1B (Fig. 1C, H and Supplemental Fig. 1E). This suggests that the influence of PRC1 on chromosomal topology extends beyond just those sites immediately bound by polycomb proteins and could explain the increase in nuclear size in the absence of RING1B (**Fig. 1A**).

### PRC1-Mediated Interactions Have a Profound Influence on Gross Nuclear Organisation

To investigate the possibility that PRC1-mediated interactions influence the topology of intervening chromatin, we interrogated the spatial proximity of additional genomic sites that lacked detectible H3K27me3 and RING1B signal. We performed 3D-FISH using pairs of fosmid probes spaced 300 kb apart and located approximately 0.5 Mb from the nearest RING1B peak (Supplemental Fig. 2A). In line with our previous observation, two independent loci showed a significant increase in inter-probe distances in *Ring1b^-/-^* mESCs compared with *Ring1b^+/+^*controls (**Fig. 1G**, **Supplemental Fig. 1E** and **Supplemental Fig. 2A**). However, we noted no such effect for a negative probe pair on chromosome 6 (**Fig. 1F** and **Supplemental Fig. 1D**). Consequently, analysis of single sites in this manner provided only limited insight into the impact of RING1B loss on PRC1 bound and non-bound chromatin.

To address this, we used oligonucleotide probe pools to investigate more extensively the impact of RING1B loss on the 3D arrangement of loci with and without PRC1 occupancy. We designed two sets of probe pools covering 28 discrete loci along a 51 Mb portion of chromosome 2. The polycomb positive probe (PcG+) was targeted to H3K27me3 ‘peaks’ enriched for RING1B occupancy (**Supplemental Table 1** and **Fig. 2A** - red bars). The non-polycomb probe (PcG-) was designed against non-repetitive sites lacking detectible H3K27me3 or RING1B ChIP-seq signal and was offset in the linear genome relative to the regions covered by the PcG+ probe (**Supplemental Table 1** and **Fig. 2A** – black bars). 3D-FISH of *Ring1b^+/+^*mESCs co-hybridised with these two probe-sets showed that the PcG-sites were significantly less clustered than those marked by the PcG+ probe (Fig. 2B, C and **Supplemental Fig. 2B**). As expected, loss of RING1B led to a significant loss of clustering between PcG+ loci but also a, albeit more subtle, reduction in the clustering of the intervening non-polycomb sites (Fig. 2B, C and **Supplemental Fig. 2B**). Compatible with our earlier observations, this suggested that reduced PRC1 levels in *Ring1b^-/-^*mESCs impacts the conformation, not only of those sites directly associated with polycomb, but of the intervening chromatin also.

**Figure 2.**
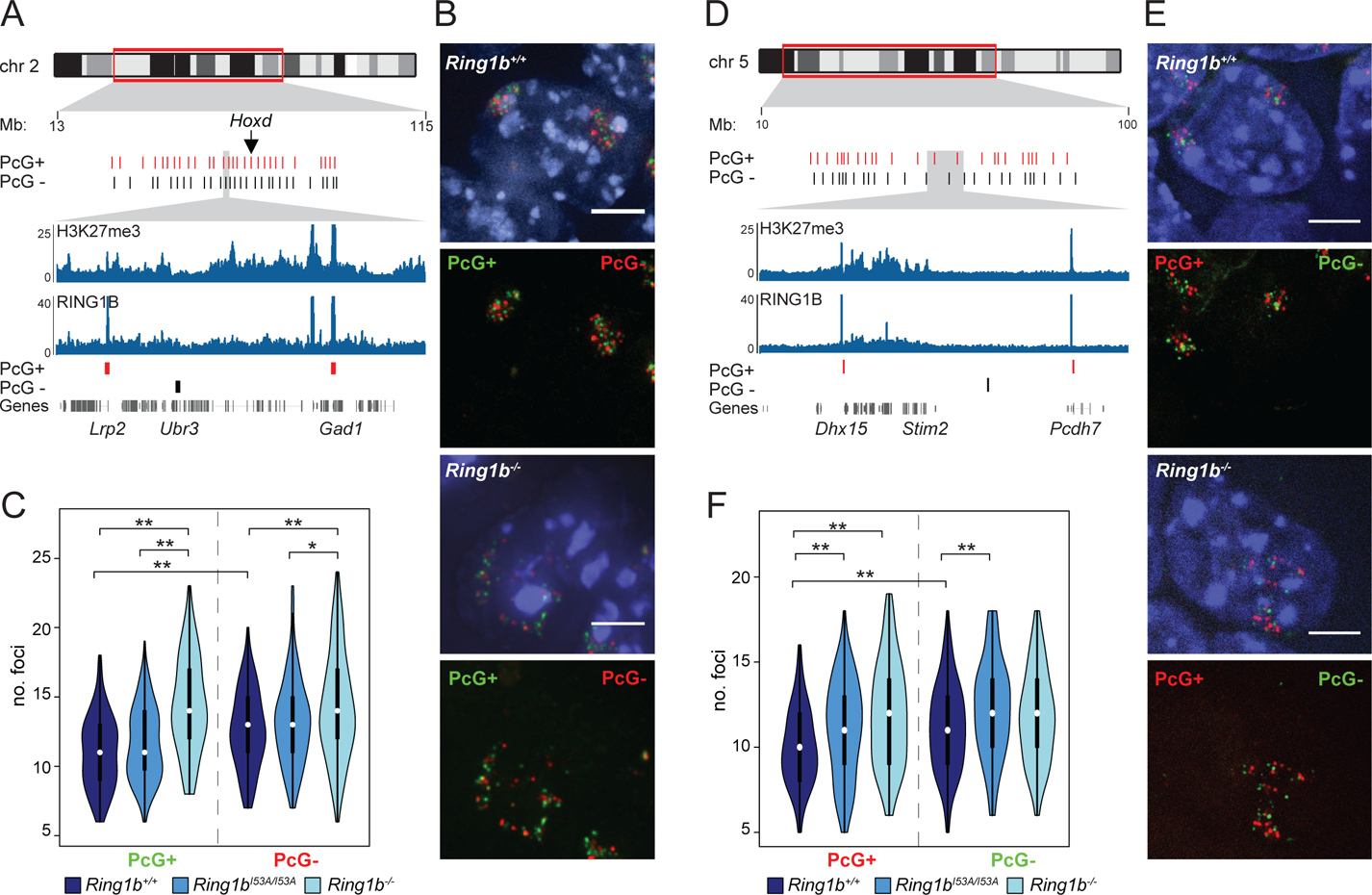
Loss of PRC1 reduces nuclear clustering at sites with and without RING1B. (A) Ideogram of chromosome 2 indicating the location of the oligonucleotide probes used in (B) and **(C)** and zoomed in browser tracks of RING1B and H3K27me3 ChIP-seq from wildtype mESCs (Illingworth et al. 2015). Polycomb positive (PcG+) and negative (PcG-) are represented as red and black bars respectively. Genomic locations are for the mm9 genome assembly. (**B**) Representative 3D FISH images of *Ring1b^+/+^* and *Ring1b^-/-^* mESCs hybridised with the chromosome 2 oligonucleotide PcG+ (green; 6FAM) and PcG-(red; ATT0594) probes. Scale bar = 5 μm. (**C**) Violin plots depicting the number of discrete foci in *Ring1b^+/+^*, *Ring1b^I53A/I53A^* and *Ring1b^-/-^* mESCs detected by the PcG+ and PcG-FISH probe pools. *p ≤ 0.05 and **p ≤ 0.01; Mann Whitney test. (**D-F**) As for A-C but for a second set of oligonucleotide probes targeted to chromosome 5. (**E**) PcG+ (red; ATT0594) and PcG-(green; 6FAM).

Strikingly, mESCs carrying a homozygous mutation encoding an isoleucine to alanine substitution at amino acid 53 of RING1B (*Ring1b^I53A/I53A^*) that profoundly impairs its E3 ubiquitin ligase activity (Buchwald et al. 2006; Eskeland et al. 2010; Endoh et al. 2012; Illingworth et al. 2015; Pengelly et al. 2015) did not disrupt clustering between either of the sets of loci being interrogated (**Fig. 2C**). This suggests that, as for local chromatin compaction (Eskeland et al. 2010) the catalytic activity of RING1B does not directly contribute to its ability to alter chromatin architecture and supports the notion that chromatin architecture and histone ubiquitination are functionally separable (Isono et al. 2013; Taherbhoy et al. 2015; Rose et al. 2016; Kundu et al. 2017; Fursova et al. 2019). However recent insights have highlighted the importance of ncPRC1 mediated H2AK119Ub in the targeting of cPRC1 (Blackledge et al. 2014; Cooper et al. 2014; Rose et al. 2016; Fursova et al. 2019). We reasoned therefore that some genomic regions might be more sensitive to impaired RING1B catalytic activity and this might have a secondary influence on chromatin architecture and nuclear organisation. To investigate this we designed another oligonucleotide probe set to target a region which displayed a more pronounced loss of RING1B binding in *Ring1b^I53A/I53A^*mESCs (**Supplemental Fig. 2C**)(Illingworth et al. 2015). These probe-sets spanned approximately 60 Mb of chromosome 5 and covered 25 discrete loci (**Supplemental Table 1** and **Fig. 2D**). As for chromosome 2, the PcG+ sites were significantly more clustered than a set of intervening sequences that lacked discernable polycomb signal (H3K27me3 and RING1B; Fig. 2E, F and **Supplemental Fig. 2D**).

Polycomb positive and negative sites also showed reduced clustering in cells lacking RING1B but, as for chromosome 2, the effect on the PcG-sites was more subtle with only one of the two experiments yielding a significant reduction in clustering (**Fig. 2F** and **Supplemental Fig. 2D**, p = 0.09 and 0.04 respectively). Interestingly, unlike chromosome 2, clustering of both PcG+ and PcG-sites within this region of chromosome 5 was substantially impaired in the catalytically deficient RING1B mutant (*Ring1b^I53A/I53A^*) mESCs in line with a more pronounced reduction in RING1B occupancy in this region in these cells (**Supplemental Fig. 2C** and **Fig. 2F**).

Taken together these findings suggest that PRC1 influences gross nuclear organisation by altering not only the polycomb bound portion of the genome but also by affecting the conformation of intervening chromatin. Furthermore, RING1B mediated ubiquitination does not directly contribute to polycomb-dependent nuclear clustering, yet its loss indirectly disrupts the association of sites where RING1B binding is substantially reduced.

### Genome-wide Compaction of Polycomb Targets

To investigate what influence PRC1 binding has on genome-wide 3D organization, we performed *in situ* Hi-C on *Ring1b^+/+^*, *Ring1b^I53A/I53A^*, and *Ring1b^-/-^* mESCs to obtain a total of ∼300 million contacts longer than 1 kb. Given the dramatic changes in nuclear size, and chromatin organization observed by FISH, we were surprised to find that the dependency of contact probability on genomic distance was largely unaffected by *Ring1b* mutations (**Supplemental Fig. 3A**). Next we analysed the impact of *Ring1b* mutations on A/B-compartmentalization using eigenvector decomposition. At this scale (200 kb), *Ring1b^+/+^* and *Ring1b^I53A/I53A^* were highly similar, while *Ring1b^-/-^* mESCs showed a more distinct 3D genome organization, both by clustering and principal component analysis (PCA; **Supplemental Fig. 3B**, C). A and B compartmentalisation reflects the spatial segregation of active and inactive chromatin in the nucleus, therefore this result likely reflects the more pronounced transcriptional changes in *Ring1b^-/-^* compared to *Ring1b^I53A/I53A^*mESCs (Illingworth et al. 2015).

5C analysis of candidate loci has demonstrated that chromosomal regions with high local PRC1 occupancy are folded into a discrete self-interacting configuration (Noordermeer et al. 2011; Williamson et al. 2014; Kundu et al. 2017). To investigate local compaction in our Hi-C data we first focused on the *Hox* loci; the most extended polycomb-associated loci in the mouse genome. These regions have previously been shown to be highly compacted by polycomb complexes, and to lose this compaction upon loss of PRC1 (Eskeland et al. 2010; Williamson et al. 2014; Kundu et al. 2017). Consistent with those observations, we detect regions of high interaction frequency inside the *Hox* clusters, that are completely lost in *Ring1b^-/-^*mESCs (Figure 3A, B and **Supplemental Fig. 3D**). *Ring1b^I53A/I53A^*mESCs did not display a significant loss of interactions relative to *Ring1b^+/+^*mESCs (Figure 3A, B and **Supplemental Fig. 3D**) suggesting only a modest disruption of PRC1-dependent 3D organization in cells bearing this mutation, consistent with our FISH data.

**Figure 3.**
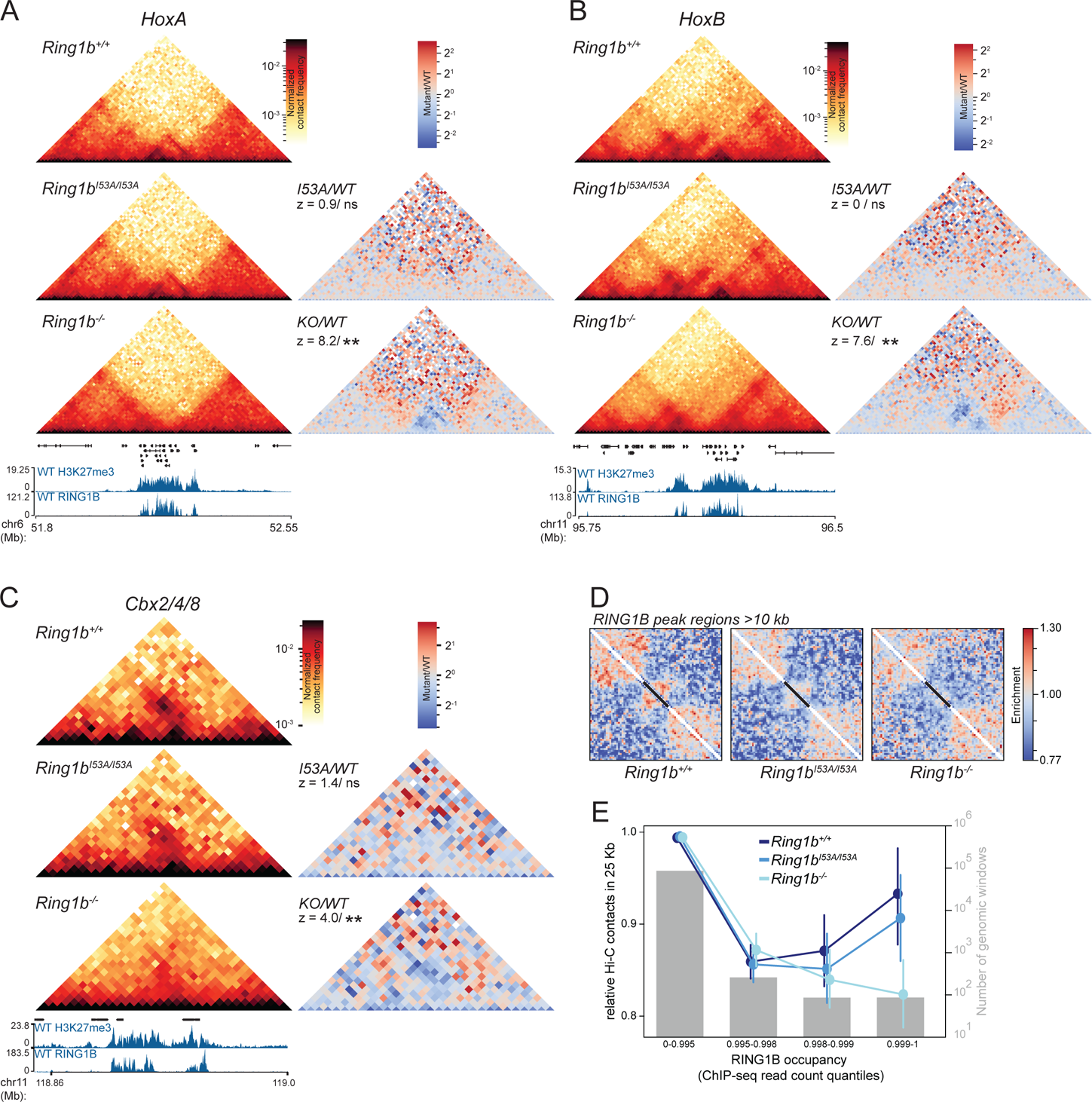
Local Compaction of PRC1 Targets. (A) Hi-C maps of the *HoxA* cluster from *Ring1b^+/+^, Ring1b^I53A/I53A^*and *Ring1b^-/-^* mESCs at 10 kb resolution. Ratios of maps from mutant over wild-type cells also shown on the right together with statistical estimation (z-score and significance level; ^ns^p > 0.05 and **p ≤ 0.01) of the difference in contact frequency within the *HoxA* cluster (statistical estimation performed on chr6 52.1 – 52.22 Mb mm9 genome build) between the cell lines. Genes, H3K27me3 and RING1B ChIP-seq profiles are shown below. **(B)** Same as **(A)**, but for *HoxB* (statistical estimation performed on chr1196.11 – 96.22 Mb). (**C**) As in (**A**), but for *Cbx2/4/8* (statistical estimation performed on chr11 118.88 – 118.96 Mb; mm9 genome build). **(D)** Re-scaled observed/expected pileups of all RING1B peak regions ≥10 kb in length (n =181) in *Ring1b^+/+^, Ring1b^I53A/I53A^* and *Ring1b^-/-^* cells. Black bars represent the location of the RING1B peak regions in the averaged map. **(E)** Average (±95% confidence interval) level of observed/expected contacts within 25 kb genomic windows split by percentiles of wildtype RING1B ChIP-seq signal shown for each of *Ring1b^+/+^, Ring1b^I53A/I53A^* and *Ring1b^-/-^*mESCs. Grey bars show number of windows in each category (right *y* axis).

To investigate if this phenomenon was restricted to *Hox* clusters, or was a more general property of extended regions of polycomb binding, we inspected other regions of pronounced polycomb association, including the *Cbx2(4/8)* and *Nr2f2* loci (**Fig. 3C** and **Supplemental Fig. 3E**). While these regions are substantially shorter than the *Hox* clusters, we observed a similar structural architecture with high local contact frequency in both *Ring1b^+/+^* and *Ring1b^I53A/I53A^* mESCs which was lost in *Ring1b^-/-^* cells (**Fig. 3C** and **Supplemental Fig. 3E**). To investigate PRC1-mediated compaction genome-wide, we performed local pile-up analysis of all 181 extended RING1B ChIP-seq peak regions (≥ 10 kb) using 1 kb resolution Hi-C data. We rescaled each region of the Hi-C map to the same length and compared the resulting pileups between each of the three mESC cell lines. Consistent with our candidate analysis, *Ring1b^+/+^*cells demonstrated a clear enrichment of local interactions corresponding to the extent of RING1B occupancy (**Fig. 3D**). This enrichment was subtly lower in *Ring1B^I53A/I53A^* cells, and completely absent in the Ring1B^-/-^cells (**Fig. 3D**). As an alternative validation of this result, we grouped all 25 kb genomic windows into quantiles of RING1B occupancy (ChIP-seq) and compared the mean observed/expected Hi-C contacts for each of these groups in each of the three mESC lines. Genomic windows bearing the highest RING1B occupancy had a local contact frequency that was substantially higher in *Ring1b^+/+^* cells than in *Ring1b^-/-^*, but *Ring1b^I53A/I53A^* were only mildly affected in line with our own and published observation (Kundu et al. 2017) (**Fig. 3E** and **Supplemental Fig. 3F**). These findings suggest that local interaction domains are a characteristic property of extended chromosomal regions bearing high levels of PRC1 binding and that this is largely independent of the catalytic activity of PRC1.

### High Levels of Canonical PRC1 Drive Distal Interactions Independently of CTCF

Beyond the scale of local interaction domains, chromatin conformation capture assays have demonstrated that distal polycomb target sites can interact and loop together into close spatial proximity (Joshi et al. 2015; Schoenfelder et al. 2015; Vieux-Rochas et al. 2015; Bonev et al. 2017; Kundu et al. 2017; McLaughlin et al. 2019). However, only a relatively small fraction of RING1B bound loci detectablly interact in our Hi-C data. Hi-C interactions between distal PRC1 binding sites were evident in our data and, as for local compaction, were largely preserved in *Ring1b^I53A/I53A^*mESCs but completely lost in cells lacking RING1B (**Fig. 4A**, Supplemental Fig. 4 A, B). To validate the interactions observed between *Bmi1* and *Skida1* (∼600 kb) we performed 4-colour FISH with three fosmid probes targetting both of these PcG+ gene loci and the intervening PcG-midpoint (Fig. 4A, B and **Supplemental Fig. 4C**). There was a significant increase in the separation between *Bmi1* and *Skida1* in both *Ring1b^-/-^*and *Ring1b^I53A/I53A^* mESCs, albeit with the later displaying a more subtle effect consistent with that observed in the Hi-C data (Fig. 4A, B). Variable levels of increase were also observed in RING1B mutant ESCs when comparing the distance separating the PcG-mid-point and either of the individual PRC1 target genes. This is consistent with the idea that PRC1 binding can impact on chromatin structure of neighbouring areas with low or undetectible levels of polycomb (**Fig. 4B** and **Supplemental Fig. 4C**).

**Figure 4.**
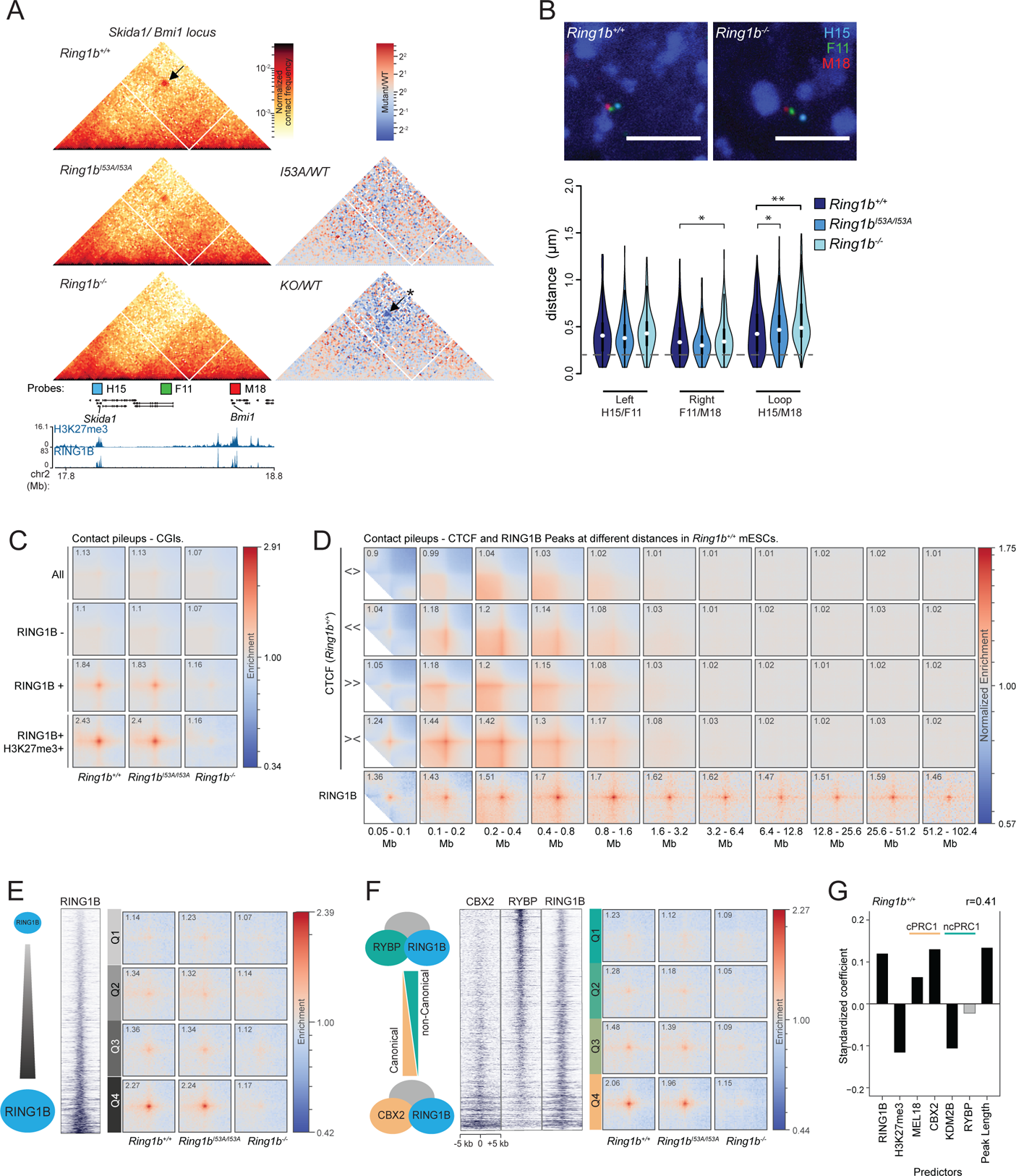
Characterisation of Distal Interactions Between PRC1 targets. (A) Hi-C data for the region of chromosome 2 harboring the *Skida1* and *Bmi1* polycomb targets. Data presented as for **Fig. 3A**. Distal interactions are highlighted with arrows and the significance of differntial signal between *Ring1b^+/+^* and *Ring1b^-/-^* Hi-C data is indicated (*p ≤ 0.05 & > 0.01). Also shown are the locations of FISH probes for the PcG+ *Skida* (H15) and *Bmi* (M18) loci and an intervening PcG-site (F11). (**B**) Representative images of 3D FISH from *Ring1b^+/+^*and *Ring1b^-/-^* cells with probes shown in (**A**). Below; violin plots show the inter-probe distances (µm) for probe pairs shown in (**A**) in *Ring1b^+/+^, Ring1b^I53A/I53A^* and *Ring1b^-/-^* cells. *p ≤ 0.05 & > 0.01 and **p ≤ 0.01; Mann Whitney test. Probes separated by < 0.2 μm (dashed grey line) are considered to be co-localised. **(C)** Pileups of interactions between CGIs. In rows: all CGIs, RING1B negative CGIs, RING1B positive CGIs, RING1B and H3K27me3 positive CGIs. In columns: *Ring1b^+/+^, Ring1b^I53A/I53A^* and *Ring1b^-/-^*cells. **(D)** Pileups of interactions between CTCF sites and RING1B peaks at different distance separations for *Ring1b^+/+^* cells. In rows: divergent CTCF sites, left-facing CTCF sites, right-facing CTCF sites, convergent CTCF sites and RING1B peaks. In columns: 2-fold increasing distance separation ranges from 0.05-0.1 Mb to 51.2-102.4 Mb. **(E)** Pileups for RING1B peaks with different level of RING1B binding by ChIP-seq. In rows: 4 quartiles of RING1B occupancy. In columns: *Ring1b^+/+^, Ring1b^I53A/I53A^* and *Ring1b^-/-^* cells. **(F)** Same as (**E**), but for quartiles of CBX2/RYBP ratio instead of RING1B occupancy. **(G)** Linear model coefficients for prediction of loop-ability of RING1B peaks for *Ring1b^+/+^* cells based on properties of RING1B peak regions (*x* axis). Positive values indicate positive impact on loop-ability. Light grey bars are not significant (p-value > 0.05). Pearson’s correlation coefficient of predicted vs observed values is shown on top right. Predictors associated with canonical and non-canonical PRC1 are denoted by ‘c’ and ‘nc’ respectively.

We quantifyied the level of interactions between PcG targets genome-wide using pileup analysis of our Hi-C data (Flyamer et al. 2019). Polycomb proteins are targeted to CpG islands (CGIs) in mammalian cells (Blackledge and Klose 2011; Deaton and Bird 2011) and so we first focused our analysis on these genomic features. We assessed distal interactions between all CGIs, CGIs lacking detectible RING1B, CGIs occupied by RING1B and CGIs associated with both RING1B and H3K27me3 (**Fig. 4C**). There was no prominent enrichment of interactions between all CGIs or CGIs lacking RING1B in *Ring1b^+/+^* mESCs, suggesting that neither the atypical base composition (high G+C and CpG) or factors associated with these regulatory elements were sufficient to coordinate distal interactions (**Fig. 4C**). In contrast, a high level of enrichment was observed between CGIs bound by RING1B, and this was further enhanced at RING1B and H3K27me3 double positive CGIs (**Fig. 4C**). Consistent with our previous observations, this enrichment was preserved in *Ring1b^I53A/I53A^* mESCs but lost in cells lacking RING1B (**Fig. 4C**).

To investigate the relationship between PRC1-mediated looping and interactions coordinated by CTCF, we performed pileup analysis of all RING1B ChIP-seq peak regions at different distance separations, and compared it to the interactions of CTCF binding sites based on their orientation (Bonev et al. 2017). This revealed loop-extrusion associated structures at CTCF site intersections, including prominent loops between convergent sites (**Fig. 4D**) (Rao et al. 2014; Sanborn et al. 2015; Fudenberg et al. 2016). CTCF-mediated loop intensities were highest between 100–400 kb, were largely undetectible at distances > 1.6 Mb (**Fig. 4D**) and were unaffected in either of the RING1B mutant mESC lines (**Supplemental Fig. 4D**). In contrast, enriched contact frequencies between RING1B binding sites were detected at distances up to ∼100 Mb (**Fig. 4D** and **Supplemental Fig. 4D**). This suggests that PRC1 sites can physically associate in cis over very large genomic distances through a mechanism that is distinct from that driving cohesin-mediated loop extrusion. To directly test this we investigated PRC1-mediated interactions in Hi-C data generated from mESCs bearing auxin-inducible degron-tagged CTCF (Nora et al. 2017). Associations between RING1B bound CGIs were unaffected by the loss of CTCF (‘auxin’; **Supplemental Fig. 4E**), confirming that the formation of PRC1-mediated interactions is mechanistically distinct from that required for loop extrusion consistent with other un-published observations (Rhodes et al. 2019).

To investigate whether the local abundance or composition of PRC1 was key to defining those sites that physically interact, we stratified all RING1B peaks into quartiles based on either ChIP-seq signal strength or peak length and performed pile-up analysis on each set of regions (**Fig. 4E** and **Supplemental Fig. 4F**). We observed a much greater enrichment of interactions between the highest-occupancy and longest RING1B peak regions (quartile 4 ‘Q4’) in both *Ring1b^+/+^* and *Ring1b^I53A/I53A^*mESCs (**Fig. 4E** and **Supplemental Fig. 4F**). Almost no enrichment was observed in *Ring1b^-/-^* cells even in Q4, consistent with our previous observations (**Fig. 4E**). This suggests that a high RING1B occupancy is critical for robust association between binding sites.

Published observations have suggested that cPRC1 complexes control polycomb-dependent 3D genome architecture, while ncPRC1s do not play a major role (Francis et al. 2004; Grau et al. 2011; Isono et al. 2013; Wani et al. 2016; Kundu et al. 2017; Plys et al. 2019; Tatavosian et al. 2019). To tested this hypothesis genome-wide, we repeated our previous analysis, this time subdividing RING1B peaks based on the ratio of ChIP-seq signal between canonical and non-canonical PRC1 subunits (CBX2 vs. RYBP respectively (Deaton et al. 2016; Rose et al. 2016)). Substantially higher interaction frequencies were observed for the relatively CBX2-enriched and RYBP-depleted peak-regions (**Fig. 4F**). A similar result was also obtained when we compared a different pair of canonical and non-canonical PRC1 subunits (PCGF2 and KDM2B respectively; **Supplemental Fig. 4G**; (Farcas et al. 2012; Morey et al. 2015)). These findings support a role for canonical PRC1 in mediating distal interactions. We noted however, that the level of RING1B enrichment was generally higher at sites relatively enriched for canonical PRC1, which, in light of our previous observations, could have potentially confounded our interpretation of these results (**Fig. 4F** and **Supplemental Fig. 4G**). Therefore, we required an alternative analysis that would allow us to independently determine the relative contribution of each factor in driving distal interactions between PRC1 targets. For this, we performed individual pileups for each RING1B peak region against all other RING1B peak regions on the same chromosome. The intensity values from the central pixel of these pileups was considered as a proxy for “loop-ability” for each region and used to build a linear model to predict the relative contribution of different parameters on the capacity to form loops (RING1B, H3K27me3, cPRC1 vs. ncPRC1 and peak length; **Fig. 4G**). As expected, we observed a strong positive contribution of the level of RING1B binding and of peak region length. Interestingly, H3K27me3 had a relatively negative impact, confirming that PRC1 and not PRC2 (which deposits H3K27me3) is important for mediating chromatin interactions. The subunits of cPRC1 (MEL18 and CBX2) both displayed a high positive effect on loop-ability, but subunits of ncPRC1 (KDM2B and RYBP) had no or negative impact, confirming our earlier analysis and previous reports (Kundu et al. 2017). Analysis of Hi-C data from *Ring1b^I53A/I53A^*cells or of an independent *Ring1b^+/+^*mESC dataset yielded an equivalent result, however data from *Ring1b^-/-^*cells dramatically reduced the coefficients for all predictors (**Fig. 4G** and **Supplemental Fig. 4H**, I). Interestingly, the only positive contributors of loop formation in the absence of RING1B were PCGF2 and CBX2 (**Supplemental Fig. 4H**). This suggests that the very low level of interactions found in *Ring1B^-/-^* cells is related to cPRC1, perhaps driven by complexes that instead incorporate the lowly expressed RING1A in place of RING1B.

### PRC1 Mediates Multivalent Interactions

Domains of high PRC1 occupancy have the potential to coordinate interactions with multiple target sites simultaneously, indeed visual inspection of our Hi-C data identified examples where adjacent RING1B peaks appeared to anchor multiple overlapping loop structures (**Fig. 4A** and Supplemental Fig. S4A). However, as this data is a population average, it was not possible to determine which, and how many, of these sites were able to interact simultaneously within an individual cell. To investigate this further we focussed on a ∼1.5 Mb portion of chromosome 5 that contains three genes which interact in a PRC1 dependent manner (*En2*, *Shh* and *Mnx1;* **Fig. 5A**). To investigate the 3D configuration of these loci in individual cells we performed 4 colour 3D FISH with probes targeting each of these genes (Fig. 5A, B). Analysis of inter-probe distances showed a significant increase in the separation between each pair of target genes upon the loss of RING1B, consistent with a loss of looping (**Fig. 5C** and Supplemental Fig. S5A). In contrast, no changes were observed between equivalently spaced probes targeting sites which lacked detectible RING1B binding (PcG-) within the same region (Fig. 5A, C and Supplemental Fig. S5A). All three genes are frequently found in proximity (all pairs ≤ 0.35 μm) and this clustering was significantly reduced upon the loss of RING1B (**Fig. 5D** and **Supplemental Fig. 5B**). The proximity of the intervening PcG-regions was not significantly altered between *Ring1b^+/+^* and *Ring1b^-/-^* mESCs (**Fig. 5D** and **Supplemental Fig. 5B**). These data demonstrate that PRC1 can coordinate interactions between multiple loci simultaneously.

**Figure 5.**
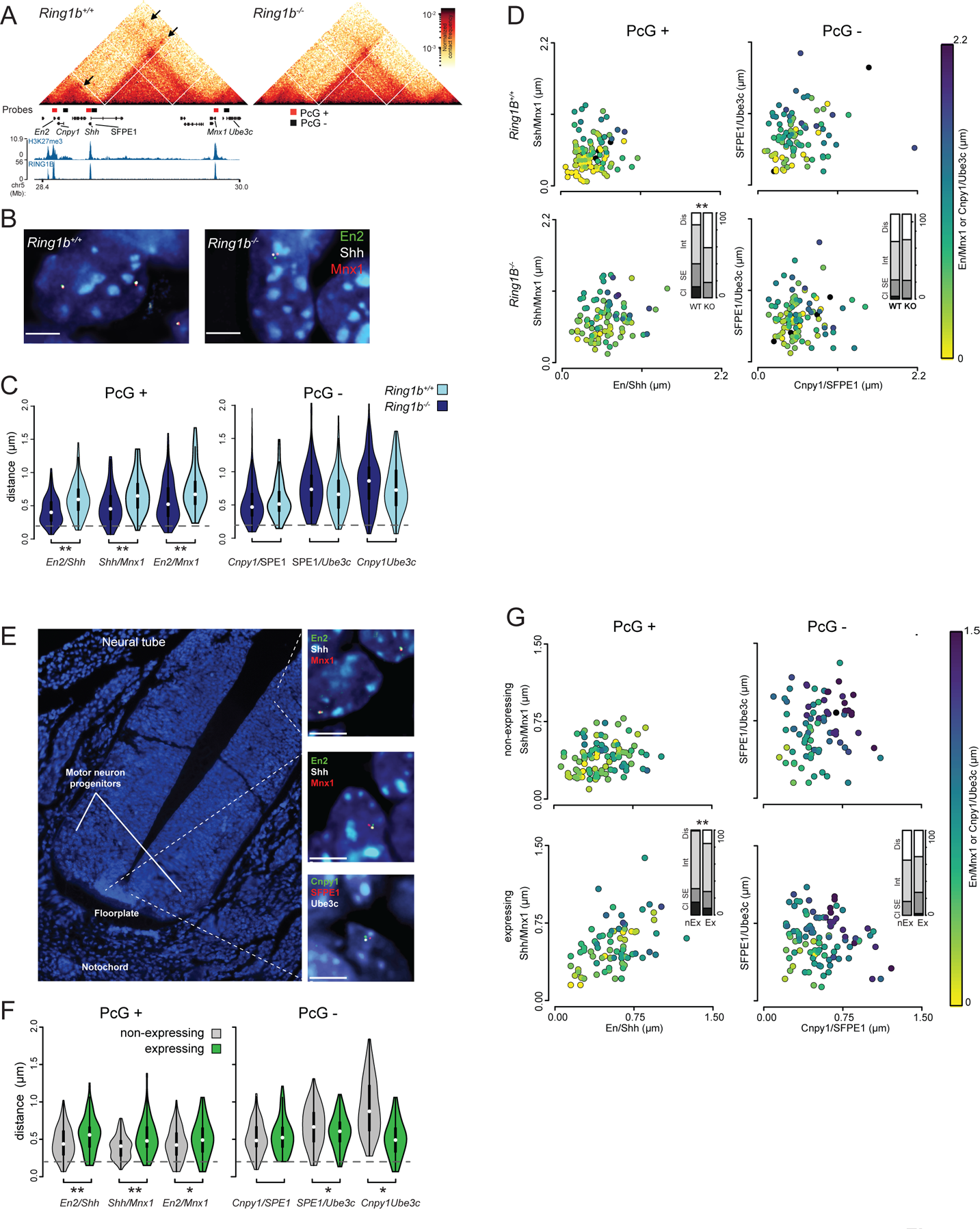
PRC1 Mediates Multivalent Interactions in Vitro and in Vivo. **(A)** Hi-C heatmaps illustrating PRC1-mediated distal interactions within the *En2 / Shh / Mnx1* locus (presented as for **Fig. 3A**; loops are highlighted with arrows). Also shown are the locations of the FISH probes used to generate (**B-G**). (**B**) Representative 3D FISH images of *Ring1b^+/+^* and *Ring1b^-/-^* mESCs hybridised with the fosmid probes shown in targeted to RING1B positive and negative sites within the locus (as illustrated in panel (**A),** red and green bars, respectively). Scale bar = 5 μm. (**C**) Violin plots depicting the distribution of inter-probe distances between each pair of fosmid probes in *Ring1b^+/+^* and *Ring1b^-/-^* mESCs. The significance of a shift in the distance between a given pair of probes between different cell lines is indicated (*p ≤ 0.05 & > 0.01 and **p ≤ 0.01; Mann Whitney test). Probes separated by less than 0.2 μm (dashed grey line) are considered to be co-localised. (**D**) Scatter plots depicting the inter-probe distances between each of the two fosmid probe pairs with the separation between the 3^rd^ pair indicated by the colour in the colour bar. Barplots representing a categorical analysis of 3-way clustering (inset) shows the percentage of nuclei which show a clustered, single excluded, intermediate or dispersed FISH signal (Cl, SE, Int and Dis respectively). A significant shift in clustering is indicated (**p ≤ 0.01; Chi-squared test). (**E**) A 3D FISH image of a transverse section through the mouse neural tube at E10.5 with zoomed inset images of 3D FISH for the fosmid probes shown in (**A**). (**F** and **G**) 3D FISH data presented as for (**C** & **D**) but for the *Shh* expressing (floorplate) and non-expressing (dorsal neural tube) cell types indicated in (**E**).

*Shh* is transcriptionally repressed and associated with RING1B and H3K27me3 in mESCs derived from the inner cell mass of the blastocyst (**Fig. 5A**). Later in development *Shh* becomes activated in a temporally and spatially restricted manner. To determine if multivalent interactions were preserved in *Shh* non-expressing cells in vivo, and if they were subsequently released upon *Shh* activation, we performed 3D-FISH on tissue sections from E10.5 mouse embryos, focussing on the floor plate and neural tube where *Shh* is expressed and repressed respectively (Delile et al. 2019). Consistent with our observation in mESCs, each individual pair of genes were more spatially separated in the expressing cells of the floor plate when compared to the cells of the dorsal neural tube where *Shh, Mnx1* and *En2* are repressed (Fig. 5E, F and **Supplemental Fig. 5C**) (Delile et al. 2019; Williamson et al. 2019). All three genes were significantly more clustered together in the dorsal neural tube than floor plate (Fig. 5F, G and **Supplemental Fig. 5D**). In contrast, the spatial arrangement of intervening control loci was not significantly different between the two regions and was substantially more dispersed in general than for the PRC1 target genes (Fig. 5F, G and **Supplemental Fig. 5C, D**). Whilst we cannot definitively conclude that *Shh*, *En2* and *Mnx1* are polycomb targets in the dorsal neural tube, they are enriched for H3K27me3 in whole neural tube tissue (**Supplemental Fig. S5E**) (Consortium 2012). We conclude that *Shh* can form multivalent interactions with other repressed genes in vitro and in vivo, and these are then subsequently lost upon gene de-repression and/or the loss of polycomb binding.

### Loss of Distal Interactions is Not Caused by Gene Activation

Compared to wild-type cells, *Ring1b^-/-^*mESCs have a more pronounced level of gene up-regulation than *Ring1b^I53A/I53A^*mESCs and have a substantially more altered chromatin structure. Moreover, we have shown that interactions between *Shh*, *Mnx1* and *En2* are lost upon gene activation. This raises the possibility that the loss of chromatin contacts in the absence of RING1B is simply a consequence of transcriptional activation. To investigate this we categorised RING1B peaks as being proximal to, or distant from, genes that are upregulated in *Ring1b^-/-^*mESCs (**Fig. 6A, B**) (Illingworth et al. 2015). RING1B peaks were classified as being upregulated if they were proximal to an up-regulated gene (situated within 0 - 50 kb from a gene with a strict definition of up-regulation -log2 expression ratio (Ring1b^-/-^/Ring1b^+/+^) ≥ 1 and a p. value of ≤ 0.01; **Fig. 6A**). Alternatively, RING1B peaks were classified as not upregulated if they were situated distant from an upregulated gene (situated > 100 kb from a gene with a liberal definition of up-regulation -log2 expression ratio (Ring1b^-/-^/Ring1b^+/+^) ≥ 0.5; **Fig. 6B**). Averaged interaction strength for each category of RING1B peak was derived from pileup analysis of Hi-C data from both *Ring1b^-/-^* and *Ring1b^+/+^* mESCs. Looping between RING1B peaks was lost in *Ring1b^-/-^* mESCs irrespective of whether the associated gene was upregulated or not (**Fig. 6A, B**), suggesting that gene activation was not responsible for the observed loss of chromatin contacts in the absence of RING1B, and similarly that loss of interactions is not sufficient to activate genes.

**Figure 6.**
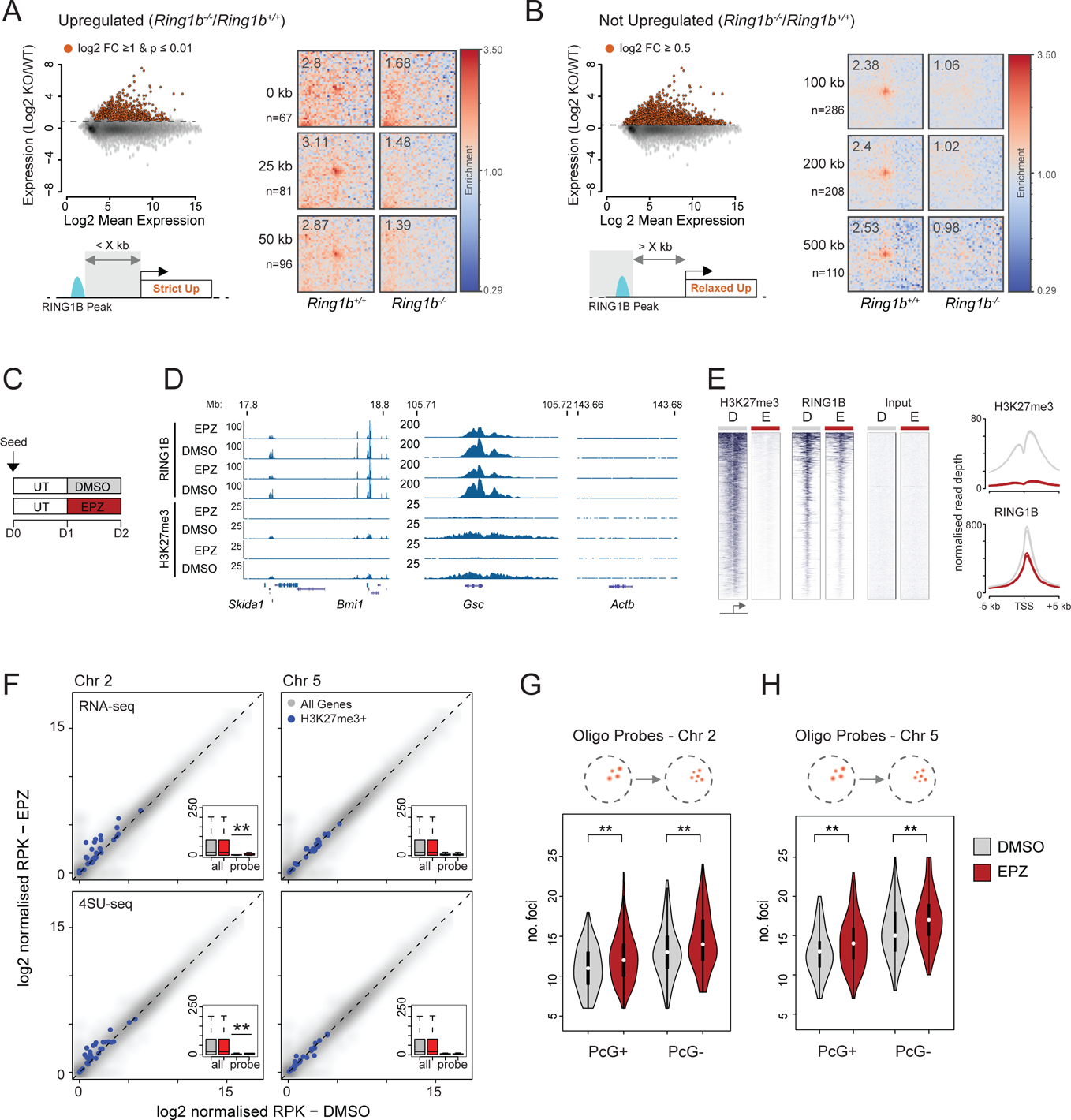
Relationship between gene upregulation and loss of PRC1-mediated looping. **(A)** Criteria on which RING1B peaks were classified as being ‘Upregulated’ in *Ring1B^-/-^* vs. *Ring1B^+/+^* mESCs. Scatter plots (upper left panel) show the log2 Mean Gene Expression versus log2 Gene Expression Ratio between the two mESC lines. Red points correspond to upregulated genes with their selection parameters noted above. Cartoon schematic (lower left panel) shows how proximity to upregulated genes (Strict Up) was used to classify RING1B peaks for subsequent Hi-C analysis. Pileup analysis of Hi-C data (right panel) illustrating PRC1 dependent distal interactions between RING1B peaks located within the indicated distance from an upregulated gene in *Ring1B^+/+^* and *Ring1B^-/-^* mESCs. n = number of RING1B peaks used in each row. (**B**) As for (**A**) but representing those RING1B peaks which are distant from upregulated genes (distant to ‘Relaxed Up’ genes). (**C**) Scheme of EPZ6438 treatment of wildtypre mESCs (UT = Serum + LIF, DMSO = Serum + LIF + DMSO and EPZ = Serum + LIF + 2.5 µM EPZ6438). (**D**) Example browser track views of RING1B and calibrated H3K27me3 ChIP-seq data from mESCs following 24 h of EPZ/DMSO treatment (2 independent replicates shown). (**E**) Heatmap representation (left panel) and summary metaplots (right panel) of RING1B and H3K27me3 ChIP-seq signal distribution at refseq gene TSS (+/-5 kb; enriched for their respective marks in wildtype ESCs) in EPZ/DMSO treated mESCs. (**F**) Scatter plots showing the relative expression (RNA-seq) or transcription (4SU-seq) levels between mESCs treated with DMSO and EPZ for 24 h. Genes located within the region covered by the oligonucleotide probes on chromosome 2 and 5 (+/-100 kb; left and right plots respectively) are highlighted in blue. Inset boxplots summarise these values for ‘all’ and ‘probe’ associated genes in the two conditions. The significance of differential expression/transcription for genes associated with the 2 probes was tested using a paired wilcoxon Rank Sum Tests and the results indicated (**p ≤ 0.01). (**G** and **H**) Violin plots depicting the number of discrete fluorescent foci in *DMSO* (grey) and *EPZ* (red), treated mESC hybridised with either the PcG+ or PcG-oligonucleotide probes on chromosome 2 (**G**) and 5 (**H**). The significance of a shift in the number of discrete foci between a given pair of samples was tested using a Mann Whitney test, the results of which are indicated (**p ≤ 0.01).

To look at a more acute response to the loss of polycomb we employed EPZ6438 (henceforth referred to as EPZ), a potent small molecule inhibitor of EZH1/2 that results in the passive loss of H3K27me3 (Knutson et al. 2013). We reasoned that treatment with EPZ for 24 h would reduce H3K27me3 levels sufficiently to reduce RING1B occupancy without substantially altering transcription (Illingworth et al. 2016). We seeded wildtype mESCs and cultured them for 24 h prior to treatment with either EPZ or DMSO (negative control) for a further 24 h (**Fig. 6C**). ChIP for H3K27me3 and RING1B followed by quantitative PCR demonstrated an almost complete loss of H3K27me3 and a substantial (approximately 50%) reduction in the level of RING1B at a panel of candidate genes (**Supplemental Fig. 6A, B**). This was confirmed genome-wide analysis by deep sequencing (ChIP-seq) (**Fig. 6D, E** and **Supplemental Fig. 6C**). We used both RNA-seq and 4SU-seq to examine the impact of EPZ treatment on total RNA and nascent transcript levels, respectively (Rabani et al. 2011). We focussed first on the *Bmi1*/*Skida1* locus (**Fig. 4A**) that showed a subtle but significant transcriptional up-regulation of the proximal genes (+/-100 kb; **Supplemental Fig. 6D, E**) and a significant increase in physical separation between *Bmi1* and *Skida1* measured by 3D FISH (**Supplemental Fig. 6F**). At the regions covered by our oligo probes on chromosomes 2 and 5 (**Fig 2A, D**), there was significant low level transcriptional upregulation across the chromosome 2 region but not for chromosome 5 (**Fig. 6F**). Despite this difference, an equivalent loss of physical clustering was observed for both oligo-probe sets (**Fig. 6G, H** and **Supplemental Fig. 6G, H**). This result supports our conclusion from Hi-C analysis that transcriptional up-regulation can be associated with, but is not required for, the loss of PRC1 mediated interactions and conversely that the loss of PRC1 mediated-interactions does not necessarily lead to gene expression, at least in ES cells.

## Discussion

Transcriptional activity, protein composition and chromatin state all play roles in specifying the spatial arrangement of the genome. Polycomb-associated facultative heterochromatin is an exemplar of this in that it mediates its own partitioning into discrete, cytologically visible, nuclear polycomb bodies (Satijn et al. 1997; Saurin et al. 1998; Pirrotta and Li 2012; Isono et al. 2013; Wani et al. 2016; Plys et al. 2019; Tatavosian et al. 2019). This organisation is established by PRC1 subunits which drive the formation of local compaction domains and longer-range chromatin interactions (Isono et al. 2013; Wani et al. 2016; Kundu et al. 2017). In this study we highlight the substantial contribution of PRC1-mediated interactions in controlling overall nuclear architecture and explore its connection to gene activity.

### PRC1 and Nuclear Architecture

Using FISH, we demonstrate that cells lacking RING1B have substantially larger nuclei and display reduced clustering of polycomb target loci. This suggests that whilst occupying less than 1% of the linear genome, RING1B has a marked impact on global nuclear organisation. By Hi-C, we observe that only those loci with the most pronounced binding of canonical PRC1, and not non-canonical PRC1, produced detectible interactions (**Fig. 4E – G** and **Supplemental Fig. 4F-I**). By comparing chromatin contacts in the presence and absence of RING1B or CTCF, we conclude that PRC1-mediated interactions are independent of CTCF, a finding consistent with other unpublished observations (Rhodes et al. 2019). PRC1 anchors chromosomal contacts at genomic distances of up to one hundred times that of CTCF, suggesting that this architecture is distinct from and independent of TADs, both in terms of mechanism and scale. We also show that PRC1-mediated chromatin interactions can be multivalent (**Fig. 5**). This is consistent with the observation that canonical PRC1 can drive liquid-liquid phase separation; a biophysical process which depends on weak multivalent interactions and which can lead to nuclear compartmentalisation and the segregation of both active and inactive chromatin states (Hnisz et al. 2017; Larson and Narlikar 2018) (Plys et al. 2019; Tatavosian et al. 2019). The polymeric nature of chromatin means that clustering of PcG sites will impose topological constraint on the intervening non-polycomb associated portion of the genome (**Fig. 7**).

**Figure 7.**
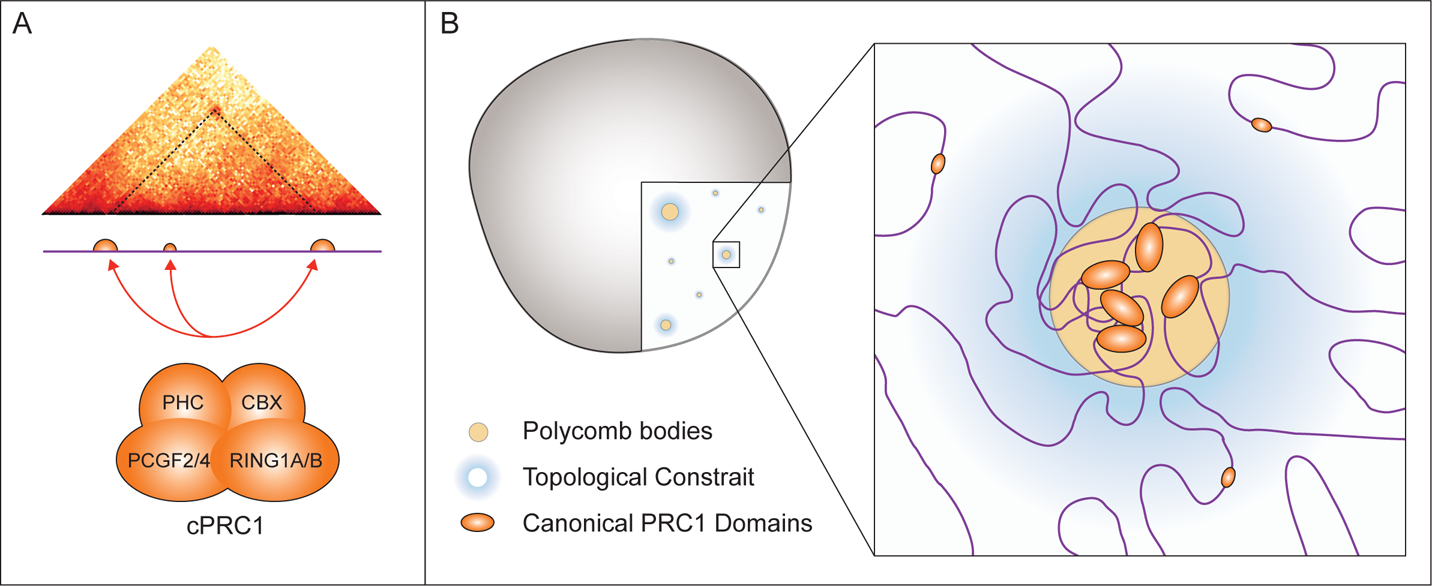
Canonical PRC1 Influences Gross Nuclear Organisation. (A) PcG mediated interactions occur preferentially between sites with the most pronounced and extended domains of canonical PRC1 occupancy. **(B)** We propose that PRC1 bound loci cluster together in the nucleus to form descrete polycomb bodies. This clustering, coupled with the polymeric nature of chromatin, imposes topological constraint on intervening non-PcG associated portion of the genome.

### PRC1 Functionality and Gene Expression

PRC1 modulates both the structure and modification state of chromatin; functions grossly ascribed to canonical and non-canocial PRC1 respectively (Grau et al. 2011; Isono et al. 2013; Blackledge et al. 2014; Rose et al. 2016; Kundu et al. 2017; Blackledge et al. 2019; Plys et al. 2019; Tatavosian et al. 2019). What then is the relative contribution of these functions to gene represion? Here we show that a hypomorphic form of RING1B, with substantially impaired catalytic activity largely preserves normal chromosomal architecture (**Fig. 2 – 4** and **Supplemental Fig. 2-4**). As this mutation yields only modest gene expression changes (Illingworth et al. 2015; Kundu et al. 2017; Cohen et al. 2018), this suggests that modulating chromosomal architecture and not H2AK119Ub is the primary repressive activity. However, induced disruption of PCGF2 and 4 which cripples canonical PRC1 specifically, leads to minimal gene up-regulation (Fursova et al. 2019). Furthermore, induced loss of variant PRC1 or a more complete disruption of RING1B E3 ubiquitin ligase activity leads to a substantial upregulation of gene expression in mESCs (Blackledge et al. 2019; Fursova et al. 2019). This suggests that low levels of H2AK119Ub are sufficient for PRC1-mediated gene repression and chromatin folding, possibly by contributing to efficient polycomb recruitment (Blackledge et al. 2014; Cooper et al. 2014).

Does the architectural function of PRC1 play any role in gene regulation? Mice bearing mutations in some subunits of canonical PRC1 display homeotic transformations indicative of mis-regulation of some Hox genes (Isono et al. 2013; Lau et al. 2017). Although, subtle in comparion to the gastrulation arrest observed in RING1B^-/-^ embryos, this observation posits two non-mutually exclusive scenarios. Firstly, that a small subset of key developmental regulators is repressed by a PRC1-mediated refractory chromatin configuration. Indeed, in the absence of E3 ubiquitin ligase activity or variant PRC1 complexes a small subset of genes (100-200), including *Hox* genes, remain repressed, (Blackledge et al. 2019; Fursova et al. 2019). A comparison between these genes and our Hi-C data showed that over half form RING1B-dependent distal interactions in mESCs (data not shown). A second possibility is that PRC1-mediated interactions in mESCs establish an architectural configuration which facilitates subsequent gene activation. Indeed, it has been shown that polycomb-dependent interactions can connect repressed gene promoters and their poised enhancers and that this is critical for correct gene activation upon neural induction (Kondo et al. 2014; Vieux-Rochas et al. 2015).

#### Concluding remarks

In this study we show that PRC1 substantially impacts on nuclear organisation and provide the first demonstrable example of expression-state dependent multivalent interactions between polycomb target sites during embryonic development. Whilst we show that it is possible to un-couple PRC1-mediated interactions from gene repression, many expression changes do accompany loss of chromatin contacts. However, no study has directly demonstrated a causal link between PRC1’s architectural function and gene repression in mammalian cells (Bantignies et al. 2003; Bantignies et al. 2011). Consequently, further work is required to fully appreciate the role of PRC1-mediated interactions in the repression and or timely activation of gene expression programmes during embryonic development.

## Materials and Methods

### Tissue Culture

Feeder-free mouse embryonic stem cells (mESCs) including E14tg2A (129/Ola; *Ring1B^+/+^*) and the derivative lines (*Ring1B* ^I53A/I53A^ and *Ring1B* ^-/-^, (Illingworth et al. 2015)) were cultured on 0.1% gelatin (Sigma - catalogue no. G1890) coated Corning flasks in GMEM BHK-21 (Gibco - catalogue no. 21710-025) supplemented with 10 % foetal calf serum (FCS; Sigma - catalogue no. F-7524), 1,000 units/ml LIF, non-essential amino acids (Gibco - catalogue no. 11140-035), sodium pyruvate (Gibco - catalogue no. 11360-039), 50 μM 2-β-mercaptoethanol (Gibco - catalogue no. 31350-010) and L-glutamine. For passaging, 60– 90% confluent ESC culture flasks were washed with PBS, incubated for 2–3 min at RT in trypsin (0.05% v/v; Gibco 25300-054), and tapped to release. Trypsin was inactivated by adding 9 volumes of ESC medium and this mixture was repeatedly pipetted to obtain a single cell suspension. ESCs were centrifuged, resuspended in ESC medium and re-plated onto gelatin coated flasks at a density of ∼4 × 10^4^ cells/cm^2^ (determined using a hemocytometer - Neubauer). For short-term EZH1/2 inhibition experiments, ESCs were plated in standard medium at 4 × 10^4^ cells/cm^2^ and cultured for 24 h. The medium was then replaced with medium supplemented with either EPZ-6438 (BioVision - catalogue no. 2383-5; reconstituted in DMSO) at a final concentration of 2.5µM or DMSO, and cultured for a further 24 h prior to harvesting or analysis.

Feeder-dependent mESCs (*Ring1B^+/+^* and *Ring1B^−/−^* (Leeb and Wutz 2007)) were plated on a layer of mitomycin C-inactivated primary embryonic fibroblasts (PEFs; derived from E12.5 mouse embryos), and grown in DMEM (Gibco - catalogue no. 41965-039) supplemented with 15% foetal calf serum, 1,000 units/ml LIF, non-essential amino acids (Sigma - catalogue no. M7145), sodium pyruvate (Sigma - catalogue no. S8636), 2-β-mercaptoethanol (Gibco - catalogue no. 31350-010) and L-glutamine. Passaging was performed as above.

For 3D FISH, mESCs were seeded onto gelatin coated Superfrost Plus microscope slides (ThermoFisher Scientific - catalogue no. J1800AMNT). 2.5×10^5^ feeder-free ESCs were seeded onto slides and cultured for 48 h prior to processing. For feeder dependent cells, PEFs were removed through 2 consecutive rounds of pre-plating (2 × 30 min in LIF-containing medium at 37°C) before plating 1×10^6^ mESCs per slide. After approximately 6 h of incubation at 37°C cells were sufficiently adherent to process for FISH.

All centrifugation steps with live cells were performed at 330 x g for 3 min at room temperature (RT). All ESC lines used in this study were routinely tested for mycoplasma.

### DNA - 3D Fluorescent *In Situ* Hybridisation (DNA - FISH)

#### Fixation

Mouse embryonic tissue sections were prepared as previously described (Morey et al. 2007). Mouse ESCs grown on slides were fixed in 4% pFA, permeabilised in PBS/0.5% Triton X, dried and then stored at −80°C prior to hybridisation. Slides were incubated in 100 μg/ml RNaseA in 2× SSC for 1 h at 37°C, washed briefly in 2× SSC, passed through an alcohol series and air-dried. Slides were incubated at 70°C for 5 min, denatured in 70% formamide/2× SSC (pH 7.5) for 40 min at 80°C, cooled in 70% ethanol on ice for 2 min and dehydrated by immersion in 90% ethanol for 2 mins and 100% ethanol for 2 min prior to air drying.

#### Hybridisation

800 ng of each fluorescently labelled oligonucleotide probe pool (2 μl; MyTags or Roche Probes) were added to (26 μl) hybridisation mix (50% formamide, 2× SSC, 1% Tween20, 10% Dextran Sulphate), denatured for 5 min at 70°C and then snap chilled on ice. 1 μg of fosmid DNA was labelled by nick translation to incorporate green-dUTP (Enzo lifesciences), AlexaFluor 594-dUTP (Invitrogen) or aminoallyl-dUTP-ATTO-647N (Jena biosciences). 100 ng of fosmid, 6 μl of Cot1 DNA and 5 μg of sonicated salmon sperm DNA was dried in a spin-vac and then re-consituted in 30 μl of hybridisation mix. Probes were then denatured for 5 min at 80°C and reannealed at 37°C for 15 mins.

Fosmid and oligonucleotide probes were hybridized to slides under a sealed coverslip overnight at 37°C. Slides were washed the next day for 4□×□3 min in 2× SSC at 45°C, 4□×□3 min in 0.1× SSC at 60°C, stained with 4,6-diaminidino-2-phenylidole (DAPI) at 50ng/ml, mounted in Vectashield (Vector) and sealed with nail varnish.

Epifluorescent images were acquired using a Photometrics Coolsnap HQ2 CCD camera and a Zeiss AxioImager A1 fluorescence microscope with a Plan Apochromat 100x 1.4NA objective, a Lumen 200W metal halide light source (Prior Scientific Instruments, Cambridge, UK) and Chroma #89014ET single excitation and emission filters(3 colour FISH) or Chroma #89000ET single excitation and emission filters (4 colour FISH) (Chroma Technology Corp., Rockingham, VT) with the excitation and emission filters installed in Prior motorised filter wheels. A piezoelectrically driven objective mount (PIFOC model P-721, Physik Instrumente GmbH & Co, Karlsruhe) was used to control movement in the z dimension. Hardware control, image capture and analysis were performed using Volocity (Perkinelmer Inc, Waltham, MA) or Nis elements (Nikon)

Images were deconvolved using a calculated point spread function with the constrained iterative algorithm of Volocity. Image analysis was carried out using the Quantitation module.

Additional information relating to all FISH probes used in this study is outlined in Supplemental Table 1.

### Calibrated ChIP Sequencing (ChIP-seq)

Trypsinised mESCs (20 x 10^6^) were washed twice in PBS. Cells were resuspended in 250 µl of PBS and fixed by the addition of an equal volume of PBS containing 2% methanol free formaldehyde (Thermo Scientific Pierce PN28906; final concentration of 1%) and incubated at RT for 10 min. Fixation was stopped by 5 min incubation with 125 mM glycine at room temperature. Fixed cells were washed in PBS and combined at this stage with 1.3×10^6^ formaldehyde fixed S2 cells (*Drosophila melanogaster* cells; for downstream calibration of ChIP-seq data). All buffers were supplemented with 1 mM DTT and 1x Protease inhibitors (Roche, 11836170001) just prior to use. Cell pellets were resuspended in lysis buffer (50 mM Tris-HCl pH 8.1, 10 mM EDTA and 20% SDS) and incubated for 10 min at 4 °C. Lysates were diluted 1:10 in ChIP Dilution Buffer (1% Triton X-100, 2 mM EDTA, 150 mM NaCl, 2 0mM and Tris-HCl pH8.1) and sonicated, first with a single 30s pulse with a probe sonicator (labtech Soniprep 150) on ice followed by a further 45 cycles using a cooled Bioruptor (Diagenode; 1 min cycles of 30sec on / 30 sec off on ‘high’ setting at 4 °C). The sonicated extract was pre-cleared by centrifugation at 16000 g for 10 min at 4 °C. The supernatant was transferred to a fresh tube and supplemented with BSA to a final concentration of 25 mg/ml. A sample of the chromatin was retained as an input reference. Antibodies were pre-coupled to a protein A Dynabeads (Life Technologies; catalogue no. 10001D) at a ratio of 1 mg antibody per 30 ml of dynabead suspension by rotation at 4 °C for 1 h. 12 x 10^6^ and 6 x 10^6^ cell equivalents of lysate were added to 7.5 µg of anti-Ring1B (Cell Signalling D22F2) or 5µg anti-H3K27me3 (Cell Signalling C36B11) respectively and incubated for 6 h on a rotating wheel at 4 °C. Following incubation, bead-associated immune complexes were washed sequentially with ChIP dilution buffer, wash buffer A and wash buffer B, each for 10 min at 4°C on a rotating wheel followed by two washes in TE buffer at RT (wash buffer A −1% Triton X-100, 0.1% Sodium-Deoxycolate, 0.1% SDS, 1mM EDTA, 500 mM NaCl, 50 mM HEPES pH7.9; wash buffer B – 0.5% NP40, 0.5% Sodium-Deoxycolate, 1 mM EDTA, 250 mM LiCl, and 20 mM Tris-HCl pH8.1). Chromatin was released by incubating the beads in 100 µl of elution buffer (0.1 M NaHCO3 and 1 % SDS) for 15 min at 37 °C, followed by the addition of 50 µg of RNaseA and 6µl of 2 M Tris pH6.8 and incubation at 65 °C for 2 h and finally by the addition of 50 µg of proteinase K and incubation at 65 °C for 8 h to degrade proteins and reverse the cross-links. Dynabeads were removed using a magnetic rack and the chromatin purified using PCR Purification columns (Qiagen) according to manufacturer’s instructions.

Libraries were constructed using the NEBNext® Ultra™ II DNA Library Prep Kit for Illumina according to manufacturer’s instructions (NEB - catalogue no. E7645S). Library PCRs were supplemented with 2x SYBR dye (Sigma – catalogue no. S9430) so that amplification could be monitored by quantitative PCR on a Roche lightcycler 480. To allow for sample multiplexing, PCRs were performed using index primers (NEBNext Multiplex Oligos for Illumina - Set 1. Catalogue no. E7335) and amplified to linear phase. Size selection purifications following the ligation and amplification PCR steps were performed with 1x and 0.9x reaction volumes of Agencourt AMPure XP beads (Beckman Coulter - A63880). Purified libraries were combined as a 12 sample equimolar pool containing the indexes 1-12 and sequenced on an Illumina NextSeq on a single high-output flow cell (single-end 75 bp reads).

### 4SU Sequencing (4SU-seq)

4SU-seq was performed essentially as described previously (Rabani et al. 2011). Briefly, 4-thiouridine (4SU; Sigma - catalogue no. T4509) was added to ESCs in culture to a final concentration of 500 µM and incubated at 37°C for 20 min. Cells were harvested by trypsinisation and washed twice with PBS at RT. Total RNA was isolated from 7×10^6^ cells using trizol according to the manufacturer’s instructions (Invitrogen - catalogue no. 15596026). Following precipitation, purified RNA was resuspended in 100 μl RNase-free water and DNase treated using the TURBO DNA-free kit according to the manufacturer’s instructions (Invitrogen - catalogue no. AM1907M). Residual inactivation beads were removed by spinning the RNA sample through a QIAshredder column at 1000 g for 1 min (Qiagen-catalogue no. 79654). 2μg of total RNA input was retained for each sample and 30μg was incubated for 1.5 h at RT with 100 μg of Biotin-HPDP (Pierce - catalogue no. 21341; reconstituted in Dimethylformamide at 1 mg/ml) in 1x Biotinylation Buffer (10 mM Tris pH 7.4 and 1 mM EDTA) to a total volume of 500 μl. Uncoupled biotin was removed through two consecutive rounds of 1:1 v/v chloroform extraction followed by isopropanol/NaCl precipitation. RNA was resuspended in 100 μl of RNase free water and mixed 1:1 w/w with µMacs Streptavidin beads (miltenyi - catalogue no. 130-074-101) and incubated for 15 min at RT with rotation. The RNA / bead mixture was applied to a µMacs column following pre-equilibration with wash buffer (100 mM Tris pH 7.5, 10 mM EDTA, 1 M NaCl and 0.1% Tween20). The captured beads were then washed with 3 x 900 µl of 65 °C wash buffer and 3 x 900 µl RT wash buffer. RNA was then eluted from the column by adding two consecutive rounds of 100 mM DTT. The eluate was added to 700 µl Buffer RLT (RNeasy MinElute Cleanup Kit; Qiagen catalogue no. 74204) and then purified according to the manufacturer’s instructions. Prior to library preparation, ribosomal RNA was depleted from both the total and purified nascent RNA using the low Input RiboMinus Eukaryote System v2 kit according to the manufacturer’s instructions (Ambion - catalogue no. A15027).

Libraries were constructed using the NEBNext® Ultra™ II Directional RNA Library Prep Kit for Illumina according to the protocol for ribosome depleted RNA and with a 11 min RNA fragmentation step (NEB - catalogue no. E7760). Library PCRs were supplemented with 2x SYBR dye (Sigma – catalogue no. S9430) so that amplification could be monitored by quantitative PCR on a Roche lightcycler 480. To allow for sample multiplexing, PCRs were performed using index primers (NEBNext Multiplex Oligos for Illumina - Set 1. Catalogue no. E7335) and amplified to linear phase. Size selection purifications following the ligation and amplification PCR steps were performed with 1x and 0.9x reaction volumes of Agencourt AMPure XP beads (Beckman Coulter - A63880). Purified libraries were combined as an 8 sample equimolar pool containing the indexes 5-12 and sequenced on an Illumina NextSeq on a single high-output flow cell (paired-end 75 bp reads).

### ChIP-seq analysis

Published ChIP-seq data from mESCs (GEO accessions: RING1B - GSM1713906-7; H3K27me3 - GSM1713910-11; MEL18 - GSM1657387; CBX2 - GSM2080677; KDM2B - GSM1272789-91; RYBP - GSM2192980-82) (Blackledge et al. 2014; Illingworth et al. 2015; Morey et al. 2015; Deaton et al. 2016; Rose et al. 2016) was retrieved from the short read archive (SRA). SRA files was converted to fastq using *fastq-dump* from the SRA Toolkit.

#### Mapping and processing

ChIP-seq data was mapped to the mouse genome (mm9 build) using *bowtie2* with the --local --threads 3 -S options to generate SAM files. Using the HOMER package, SAM files were converted into tag directories and multi-mapping reads were removed using *makeTagDirectory* -unique -fragLength 150. Mapped regions which, due to fragment processing, extended beyond the end of the chromosomes were removed using *removeOutOfBoundsReads.pl* with chromosome lengths for mm9. Replicate data, where appropriate, was combined at this stage. Genome browser files (.bw) were generated using *makeUCSCfile* with the -bigWig -fsize 1e20 -norm 10e7 -color 25,50,200 options. H3K27me3 genome browser files were normalised instead to a calibrator value set to maintain the relative contribution of Drosophila spike-in reads between the input and immunoprecipitated samples.

#### Signal quantitation

For ChIP quantitation, published RING1B peaks (Illingworth et al. 2015) separated by less than 5000 bp were merged using bedtools *mergeBed* function with -d 5000. HOMER was then used to quantify read coverage across these merged regions. For linear modelling (see **Hi-C data analysis below**) simple read coverage was determined using a*nnotatePeaks.pl* with the following parameters -size “given” -noann -nogene -len 0 - strand both -norm 10e7. Window files centred on RING1B peaks (+/-5 kb) used to make heatmaps were generated using a*nnotatePeaks.pl* **-**size 10000 -hist 200 -ghist -nogene - strand both -norm 10e7 (calibrated normalisation for H3K27me3 chIP-seq data was performed as outlined above). For the comparison of HiC-contact frequency to RING1B/H3K27me3 occupancy, ChIP signal was quantified across the whole mouse genome in 25 kb abutting windows using *annotatePeaks.pl* with the same parameters as simple RING1B peak quantitation outlined above. Where appropriate all quantifications were expressed as reads per kb per million mapped reads (RPKM).

#### CGI analysis

The coordinates of biochemically defined mouse CGIs (mm9; (Illingworth et al. 2010)) were intersected with published peaks of RING1B and H3K27me3 (Illingworth et al. 2015) using the *intersect* function of *bedtools* with the following paramaters -wa -u –a.

### 4SU-seq & RNA-seq analysis

#### Mapping and processing

For each de-multiplexed sample, multiple raw fastq files were merged (individually for reads 1 and 2) and then aligned to the mouse genome (mm9) using bowtie2 v2.2.6 for paired end sequence data (options: --local --threads 3) to generate .SAM files. Aligned read data was processed using HOMER v4.8. SAM files were converted into tag directories using ‘makeTagDirectory’ with the following parameters: -format sam -flip –sspe. Genomic intervals which extended beyond the end of the chromosomes were removed using ‘removeOutOfBoundsReads.pl’. Strand specific browser track files (bigwig format; ‘.bigWig’) for each replicate were generated using ‘makeUCSCfile’ with the following parameters: -fsize 1e20 -strand + (or -) -norm 1e8.

#### Signal quantitation

HOMER was used to quantify 4sU/RNA-seq read coverage across all Refseq genes (mm9). Coverage was determined using a*nnotatePeaks.pl* with the parameters: -size “given” -noann -nogene -len 0 -strand both -norm 0. All expression values were then converted into reads per kb per million mapped reads (RPKM) using R (R Core Team 2019).

### In situ Hi-C

We performed *in situ* Hi-C largely in accordance with (Rao et al., 2014) with minor modifications, same as in (McLaughlin et al. 2019). Briefly, the modifications included: digestion using DpnII instead of MboI (in the DpnII buffer, with previous washes in NEBuffer 3); no phenol-chlorophorm extraction after decrosslinking with buffer exchange using Amicon filter units (30 kD, 500 µl); sonication using a probe-based sonicator to achieve fragment length distribution of ∼200-700 bp followed by concentration on Amicon filter units; indexing using barcoded primers instead of adaptors; size selection of the final amplified library through gel-extraction instead of AMPure beads. Final Hi-C libraries were test-sequenced at the Wellcome Trust Clinical Research Facility (Edinburgh) on NextSeq550 PE75, and selected high quality (by cis/trans ratio and consistent P_c_(s) curve) libraries were deep sequenced at BGI on HiSeq4000 PE 100. We had 2 (I53A and KO) and 4 (WT) replicate libraries per condition with a total of ∼0.85-1.18 billion reads.

### Hi-C data analysis

Hi-C data were analyzed using the *distiller* pipeline (https://github.com/mirnylab/distiller-nf) on Eddie3 cluster of the University of Edinburgh. Mapping was performed to the *mm9* genome assembly, PCR and optical duplicates were removed with the *max_mismatch_bp: 0* option. Data were filtered to remove reads with mapq<30, binned to generate multiresolution .cool files and balanced using default parameters. The same analysis was performed with deep Hi-C data from ES cells (Bonev et al. 2017).

For insulation analysis, we used *cooltools diamond-insulation* with 25 kbp resolution data and 1 Mbp window size. For eigenvector analysis, we used *cooltools call-compartments* with 200 kbp resolution data and reference track of GC content. For both analyses, we then clustered the genome-wide insulation profile or eigenvector using *seaborn.clustermap* with default algorithm setting.

Pile-up analysis was performed using *coolpup.py*. All distal and non-rescaled local pileups used chromosome-wide expected normalization; local rescaled pileups were normalized to randomly shifted control regions (10 per region of interest). Unless specified, we didn’t consider regions closer than 100 kb. Pileups investigating enrichment at different distance scales used *coolpup.py*’s *--mindist* and *--maxdist* options to specify distance ranges. Local rescaled pileups were created with *--rescale_size 75 --rescale_pad 2 --minsize 10000* options.

Loop-ability was calculated using *--by_window* of *coolpup.py*. We took the Enrichment1 values, corresponding to the observed/expected contacts in the central pixel of the pileups, and didn’t perform any filtering based on coefficient of variation. This table was merged with information about level of binding/occupancy of different factors determined by ChIP-seq. We only considered regions with *Enrichment1*>0 in all three datasets, and less than 1000 reads of RING1B (since regions with higher coverage represented technical artefacts). We used *scikit-learn* to perform linear modelling using a subclass of *linear_model.LinearRegression* that also calculates p-values for each predictor (https://stackoverflow.com/a/27975633/1304161). We used properties of merged RING1B ChIP-seq peaks(see above): number of reads from ChIP-seq of H3K27me3, RING1B, MEL18, CBX2, KDM2B and RYBP, and merged peak length. All predictor values were normalized using *preprocessing.StandardScaler()* method of *scikit-learn*. We then used the values of coefficients for each predictor to compare their relative importance for looping interactions between RING1B peaks.

Local compaction analysis was performed same as described in (McLaughlin et al. 2019). Briefly, total number of normalized observed/expected contacts in 25 kb windows was calculated, excluding the two first diagonals and any regions containing filtered out bins. Then this was compared to the total number of ChIP-seq reads of RING1B or H3K27me3 from these regions.

#### Expression vs. Distal Interactions

To investigate the contribution of the loss of RING1B protein versus gene de-repression on distal interactions we compared our Hi-C data to published gene expression data for *Ring1b^+/+^* and *Ring1b^-/-^* mESCs (Illingworth et al. 2015).

Refseq genes were classified as having either ‘strict’ upregulation (log2 fold change ≥ 1 and an adjusted p value of ≤ 0.01) or ‘relaxed’ upregulation (log2 fold change ≥ 0.5) in *Ring1b^-/-^* vs. *Ring1b^+/+^* mESCs. Hi-C pileups were generated for RING1B peaks with the highest ChIP-seq signal (upper quartile; Q4) either for peaks associated with upregulated gene (proximal to ‘strict’ genes) or not-associated with upregulated genes (distant from ‘relaxed’ genes). A range of gene-to-peak distances were assessed.

#### Statistical testing of differential interaction frequencies

Observed/expected signal ratios for individual genomic regions of interest (ROI) were extracted and used to determine the average level of interaction enrichment for that region for each Hi-C dataset. Matched values from 1000 random regions of the same shape and size were determined (matched for chromosome and distance from the matrix diagonal). These permuted values were subsequently used to estimate the mean and standard deviation of the distribution of all regions for the chromosome. Following log transformation these values were used to generate Z score for the region of interest. The mean was subtracted from the observed value for the ROI and divided by its standard deviation and subsequently converted into a p-value for ease of interpretation (as *1*⍰ *scipy.special.ndtr(zscore)*).

## Data Availability

All sequencing data was submitted to the GEO repository under accession numbers GSE134826 (Hi-C) and GSE140894 (4sU/RNA-seq and ChIP-seq).

## Acknowledgemenst

We are grateful to Nezha Benabdallah for help with experiments and to all members of the Bickmore lab for helpful and insightful discussions during the preparation of this manuscript. We would like to thank Elisabeth Freyer and Stacey Riddles (IGMM) for flow cytometry support and to the WTCRF (Edinburgh Clinical Research facility) for sequencing. RSI was supported by a BBSRC project grant (BBSRC_BB/H008500/1) and an MRC Career Development Award (MR/S007644/1). IMF was funded by a PhD studentship from the Darwin Trust. DS was supported by a Newton fellowship from the Royal Society. Work in the WAB lab is funded by an MRC University Unit grant MC_UU_00007/2.

---

**Figure S1.**
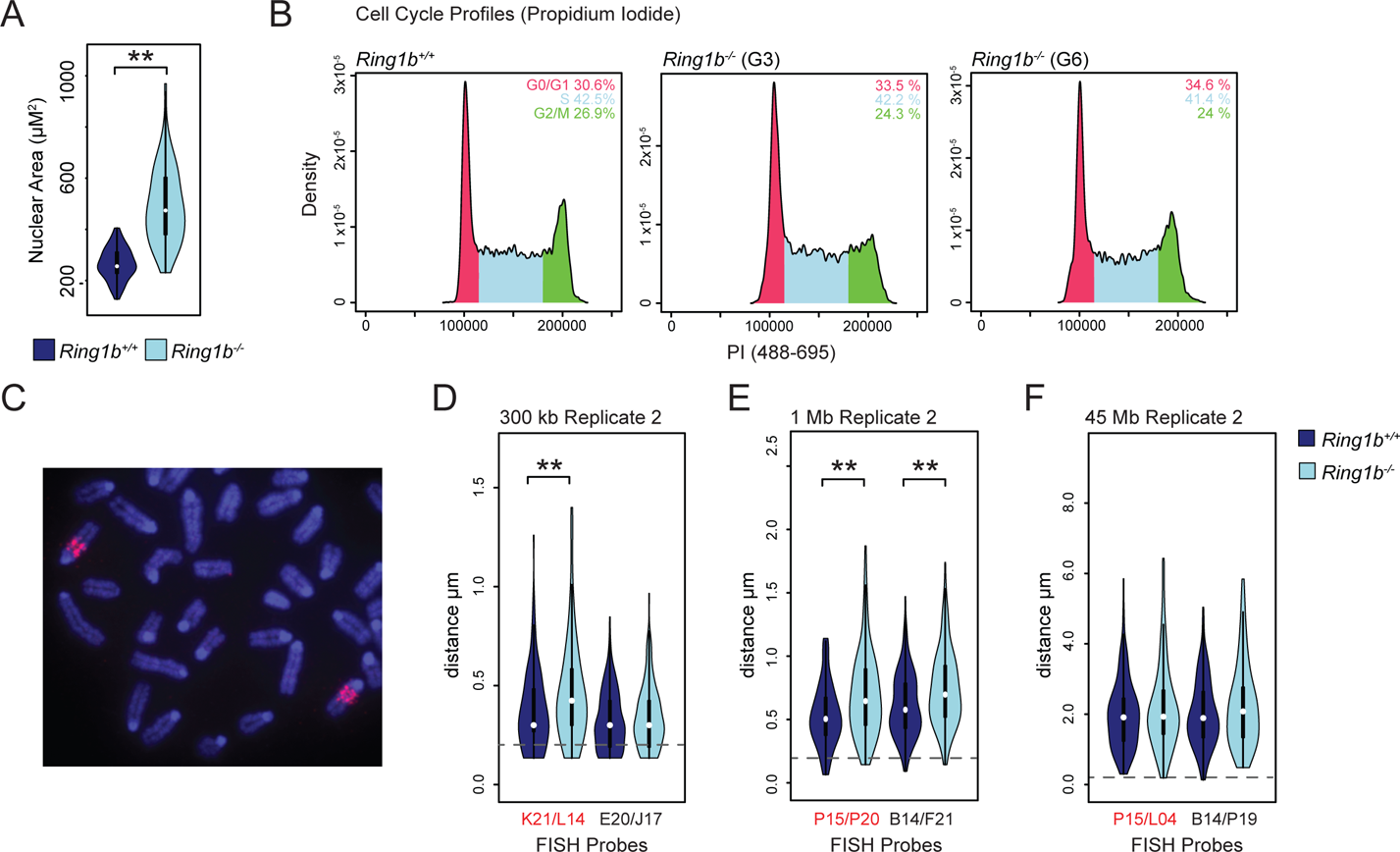
Cytological Analysis of *Ring1b^+/+^* vs. *Ring1b^-/-^*mESCs. (**A**) Violin plot depicting the nuclear area of *Ring1b^+/+^* and *Ring1b^-/-^* mESCs determined by DAPI staining of 2D nuclear preparations (independent replicate experiment to that shown in **Fig. 1A**). A significant shift in nuclear area is indicated (**p = 6.94×10^-19^; as determined by a Mann-Whitney test). (**B**) Histogram profiles of fluorescence-activated cell sorting (FACS) data from propidium iodide stained *Ring1b^+/+^* and *Ring1b^-/-^* mESCs (two independent *Ring1b^-/-^* clones are shown - G3 and G6). The relative distribution of cells in G0/G1, S and G2/M are indicated in parenthesis (red, blue and green respectively). (**C**) Representative FISH image of the chromosome 6 polycomb positive oligonucleotide probe signal on metaphase chromosomes from wildtype mESCs. (**D-F**) Violin plots of inter-probe distances for the indicated fosmids (locations shown in **Fig. 1C**) in *Ring1b^+/+^* and *Ring1b^-/-^* mESCs (independent replicate experiment to that shown in **Fig. 1G-I**). Probes separated by < 0.2 μm (dashed grey line) are considered to be co-localised. A significant shift in inter-probe distance between *Ring1b^+/+^* and *Ring1b^-/-^* mESCs is indicated (**p = 2.38×10^-3^ (**G**) and **p = 2.09×10^-5^ and **7.88×10^-5^ (**H**); as determined by Mann Whitney test).

**Figure S2.**
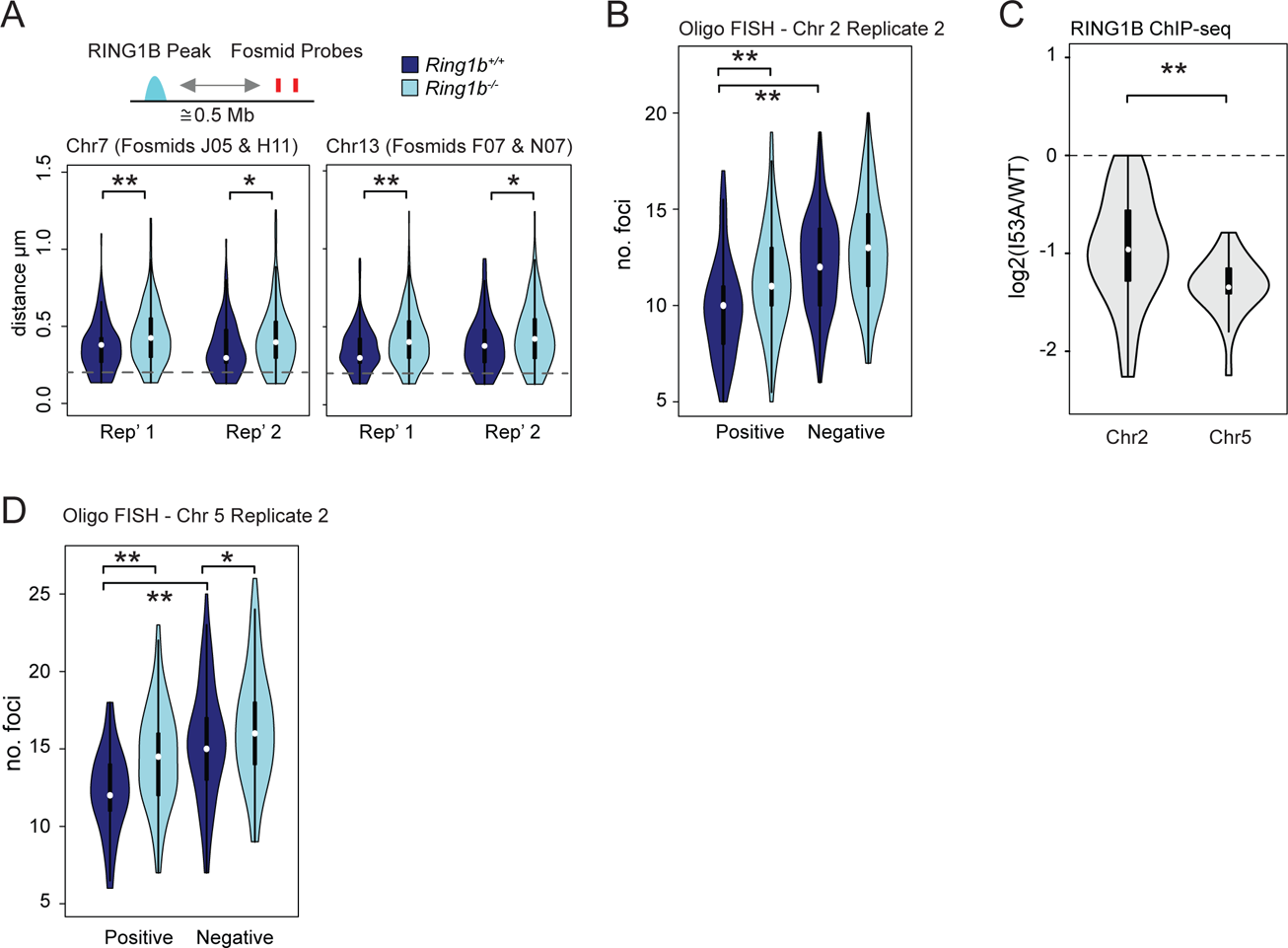
Loss of PRC1 Reduces Nuclear Clustering at Sites Lacking RING1B. (**A**) Violin plots showing the distribution of inter-probe distances determined by 3D FISH of *Ring1b^+/+^* and *Ring1b^-/-^*mESCs hybridised with fosmid probes situated at non-RING1B sites on chromosome 7 and 13 (left and right panels respectively; fosmids are located approximately 0.5 Mb from the closest RING1B peak). A significant shift in inter-probe distance, as determined by a Mann Whitney test, is indicated (**p = 7.79×10^-5^, *p = 1.53×10^-2^, **p = 6.54×10^-4^ and *p = 1.58×10^-2^ from left to right). Genomic locations are given for the mm9 genome build. Each plot shows the result of two independent replicates. Probes separated by less than 0.2 μm (dashed grey line) are considered to be co-localised. (**B**) Violin plots depicting the number of discrete fluorescent foci in *Ring1b^+/+^*and *Ring1b^-/-^* mESCs hybridised with either the polycomb positive or negative oligonucleotide probes on chromosome 2 (second independent replicate to that shown in **Fig. 2C**). The significance of a shift in the number of discrete foci between a given pair of samples was tested using a Mann Whitney test, the results of which are indicated (**p ≤ 0.01). (**C**) Violin plot showing the log2 ratio (*Ring1b^I53A/I53A^* vs. *Ring1b^+/+^*) of RING1B ChIP-seq signal across all RING1B peaks within the chromosome 2 (42,398-93,905 Mb) and chromosome 5 (42.40-93.91 Mb) regions representing the PcG+ positive and a second proposed PcG+ oligonucleotide probe respectively (± 100 kb). A significantly greater reduction in RING1B levels across the chromosome 5 region was observed (**p = 0.00114; Mann Whitney test). (**D**) Violin plots depicting the number of discrete fluorescent foci in *Ring1b^+/+^*and *Ring1b^-/-^* mESCs hybridised with either the polycomb positive or negative oligonucleotide probes on chromosome 5 (second independent replicate to that shown in **Fig. 2F**). The significance of a shift in the number of discrete foci between a given pair of samples is indicated (*p ≤ 0.05 and **p ≤ 0.01; Mann Whitney test).

**Figure S3.**
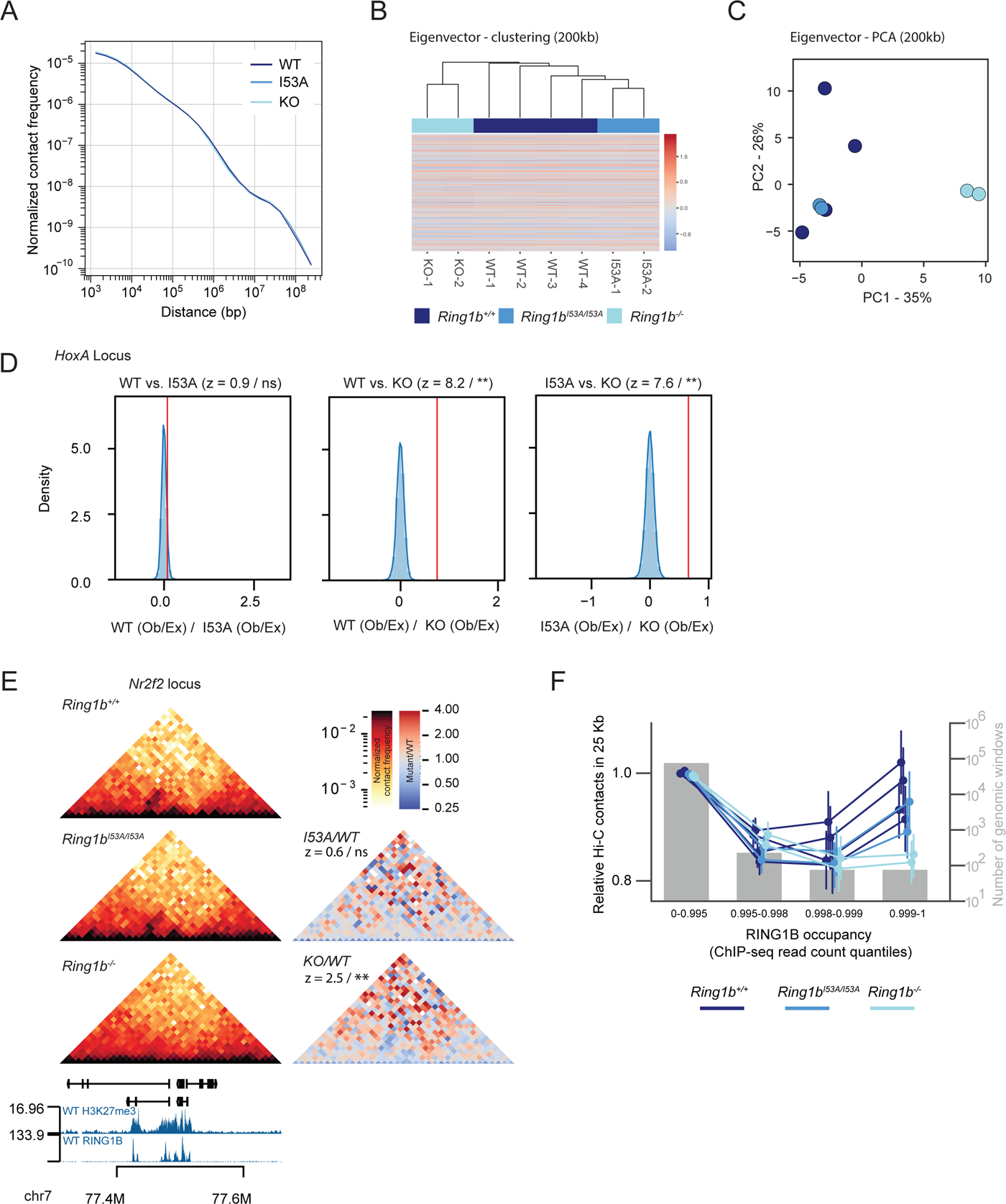
Validation of Hi-C analysis. (A) Curves of probability of contact at different distance separations for *Ring1b^+/+^, Ring1b^I53A/I53A^* and *Ring1b^-/-^* cells. **(B)** Clustering of first eigenvectors (200 kb resolution) for replicates of Hi-C data for *Ring1b^+/+^, Ring1b^I53A/I53A^*and *Ring1b^-/-^* mESCs. **(C)** PCA analysis of eigenvectors from (**B**). **(D)** Statistical estimation of contact frequency changes within the *HoxA* cluster for all pairwise comparisons between *Ring1b^+/+^, Ring1b^I53A/I53A^*and *Ring1b^-/-^* cells. Shown are histograms of ratios of observed/expected signal for 10,000 random regions on the same chromosome (bars) and its associated kernel-density estimate (curve). The red vertical line represents the value for the *HoxA* region. Z-value and level of significance shown. **(E)** Same as **Fig. 3A**, but for the *Nr2f2* region (statistical estimation performed on chr7 77.43 – 77.52 Mb; mm9 genome build). **(F)** Same as **Fig. 3E**, but for individual replicates.

**Figure S4.**
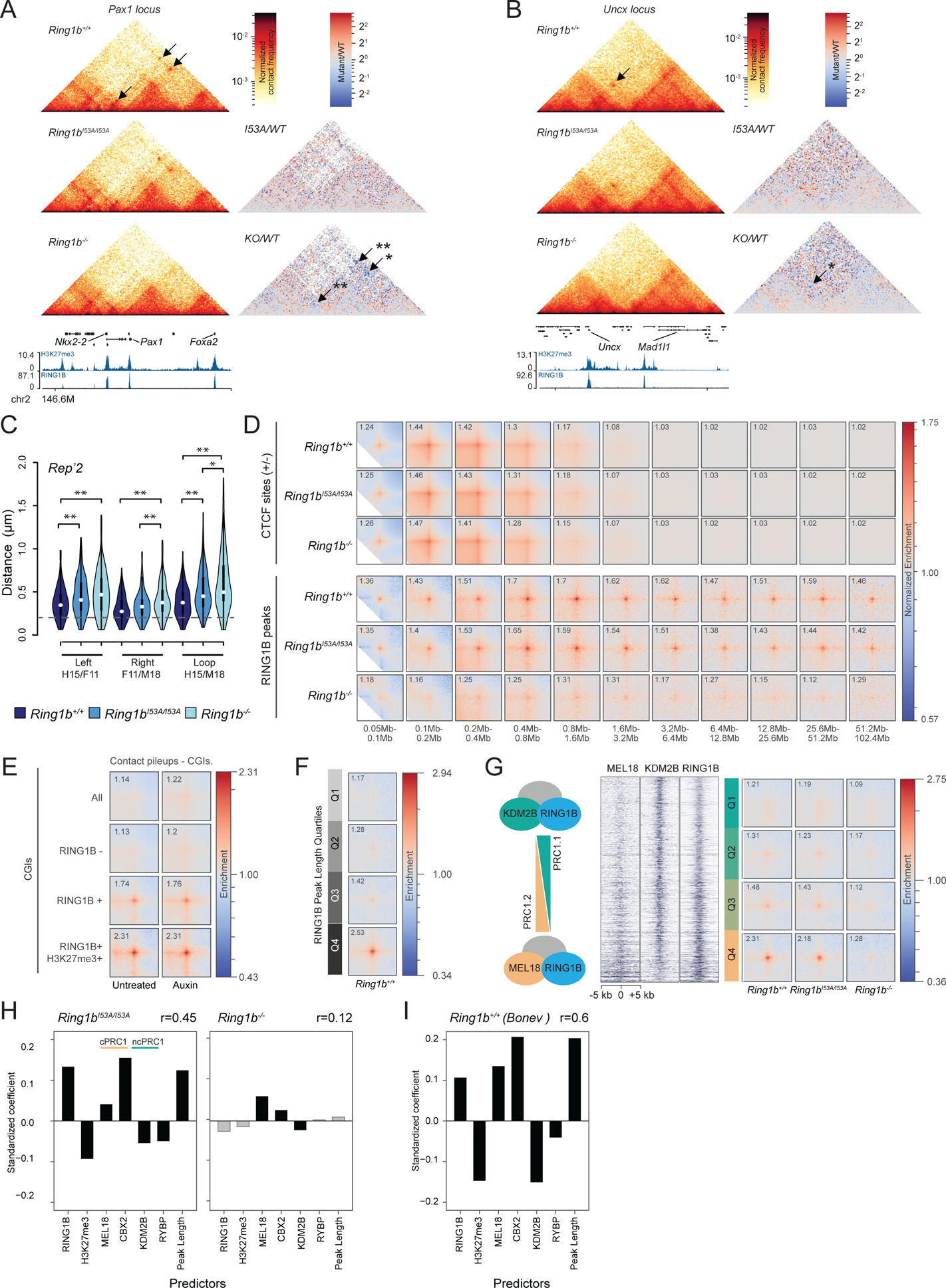
Additional Analysis of Looping Between PRC1 targets. (A) Same as **Fig. 4A**, but for the loops between RING1B peaks proximal to the *Nkx2-2*, *Pax1* and *Foxa2* genes. **(B)** Same as **Fig. 4A**, but for the loops between RING1B peaks proximal to the *Uncx* and *Elfn1* genes. **(C)** Same as Figure 4B, but for two FISH replicates separately. **(D)** Same as for **Fig. 4D**, but for *Ring1b^+/+^, Ring1b^I53A/I53A^* and *Ring1b^-/-^*cells and only convergent CTCF sites and RING1B peaks. In rows: convergent CTCF peaks for *Ring1b^+/+^, Ring1b^I53A/I53A^* and *Ring1b^-/-^* cells, then wildtype RING1B peaks in *Ring1b^+/+^, Ring1b^I53A/I53A^*and *Ring1b^-/-^* mESCs. **(E)** Same as **Fig. 4C**, but for published Hi-C data from untreated or auxin-treated CTCF-AID cells ((Nora et al. 2017)). **(F)** Same as **Fig. 4E**, but for peak region length instead of RING1B ChIP-seq signal, and only for *Ring1b^+/+^* cells. **(G)** Same as **Fig. 4F**, but for MEL18/KDM2B ratio. **(H)** Same as **Fig. 4G**, but for *Ring1b^I53A/I53A^* and *Ring1b^-/-^* cells. **(I)** Same as **Fig. 4G**, but for published data from wild-type ES cells ((Bonev et al. 2017)).

**Figure S5.**
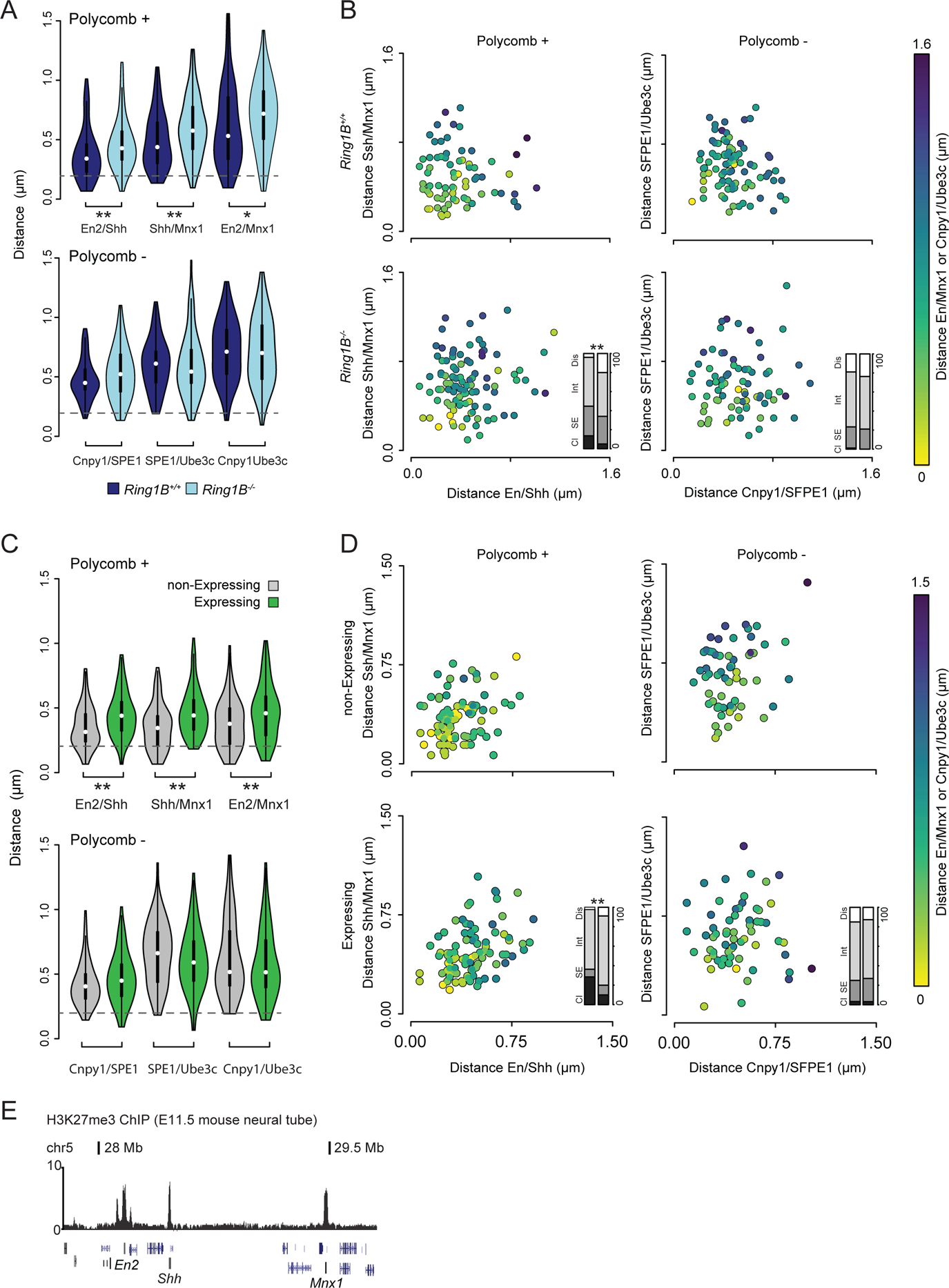
Additional Data Supporting the Existence of Multivalent PRC1-Mediated Interactions. Panels (A-D) show independent replicates of the datasets shown in **Fig. 5C**, D, F and G.

**Figure S6.**
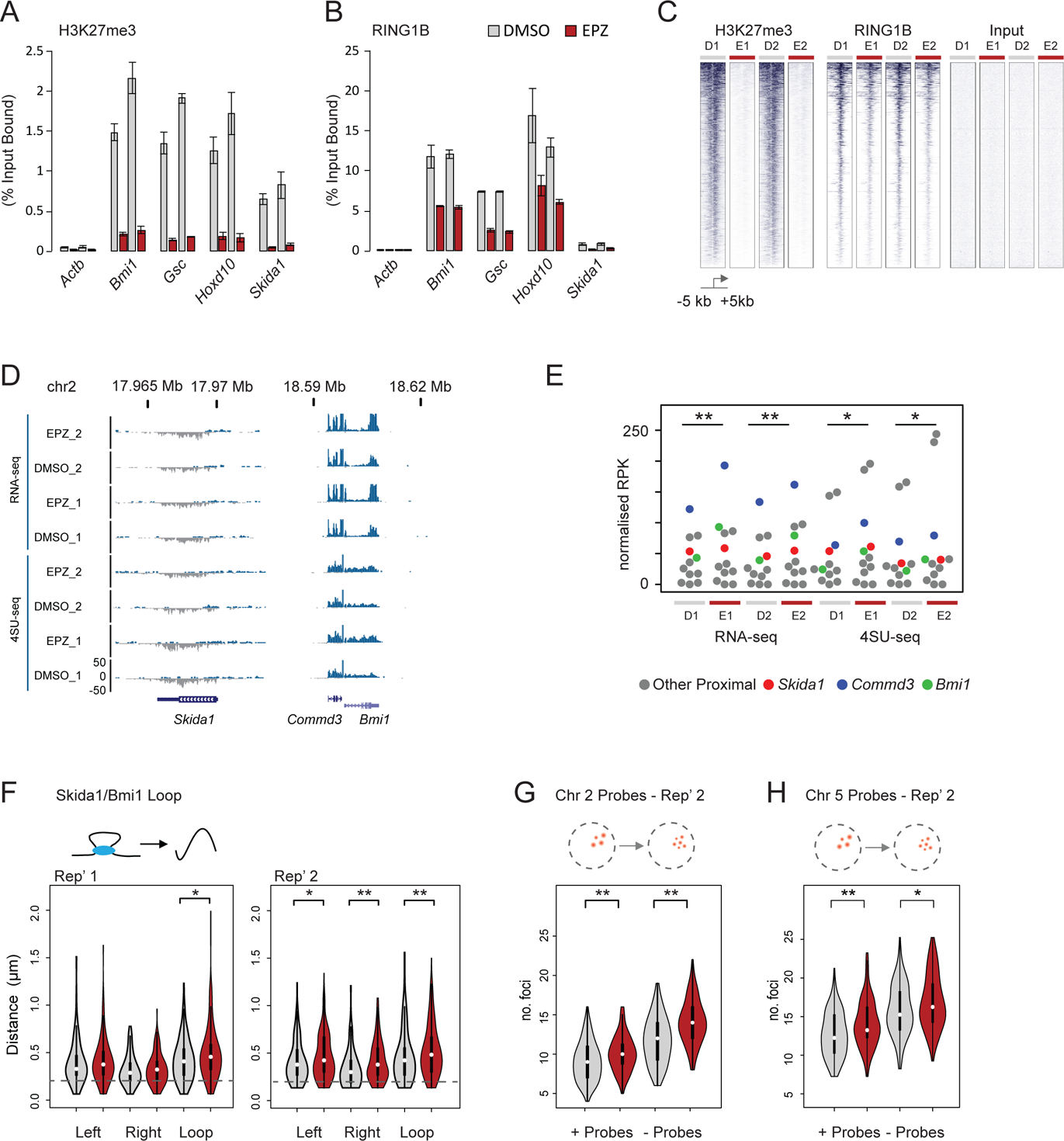
Loss of PRC1 and not Transcriptional Upregulation Results in the Loss of Chromatin Contacts. (**A & B**) Barplots representing candidate analysis of ChIP enrichment relative to input measured by quantitative PCR for H3K27me3 (**A**) and RING1B (**B**) in mESCS treated for DMSO or EPZ (two independent replicates are shown). (**C**) Heatmap representation of H3K27me3 and RING1B ChIP-seq signal distribution at refseq gene TSS (+/-5 kb; enriched for their respective marks) in two independent sets of EPZ/DMSO treated mESCs. (**D**) Example browser track views of RNA-seq and 4SU-seq data from mESCs following 24 h of EPZ/DMSO treatment (2 independent replicates shown). The signal is coloured according to the transcribed strand (positive - blue and negative -grey). (**E**) Beeswarm plots illustrating the RNA-seq and 4SU-seq signal of genes within the Skida1/ Bmi1 in mESCs treated with either DMSO or EPZ for 24h. Significant differential expression is indicated (*p ≤ 0.05 & > 0.01 and **p ≤ 0.01; paired Wilcoxon Rank Sum test). (**F**) 3D FISH measurement for probes shown in (**Fig. 4A**) in mESCs treated with DMSO or EPZ for 24h (Two independent replicate experiments are shown). The significance of a shift in inter-probe distance between a given pair of samples is indicated (*p ≤ 0.05 & > 0.01 and **p ≤ 0.01; Mann Whitney test). Probes separated by less than 0.2 μm (dashed grey line) are considered to be co-localised. (**G** and **H**) Replicated experiments for those shown in **Fig. 6G** and **H**.

